# A bistable circuit regulates miRNA-155 levels in human macrophage inflammatory transitions

**DOI:** 10.1101/2025.07.09.663909

**Authors:** Rodrigo A. Mora-Rodríguez, Jose Guevara-Coto, ManSai Acon, Jorge Torres, Guillermo Oviedo-Blanco, Anne Régnier-Vigouroux, Carsten Geiß

## Abstract

Macrophages are crucial immune regulators as they can either trigger or resolve inflammation. These properties rely on defined inflammatory states and make macrophages valuable therapeutic targets. Identification of stable steady states and bistability in inflammatory transitions provides deeper insights into immune regulation and facilitates the development of novel therapeutic strategies. We present a multiomic and systems biology approach for the top-down identification of bistable circuits in human macrophages polarized towards pro- and anti-inflammatory phenotypes. Using differential gene expression profiles, we identified three criteria to suspect bistability: two potential attractors, a hysteresis behavior between their transitions, and the presence of potential modules of coregulated genes. This was further confirmed by proteomics data pointing to mutually exclusive and time-dependent profiles of gene expression. By network simplification and creation of a novel pipeline for parameter estimation in bistable models, we obtained a minimal model of inflammatory transitions in which we identified ultrasensitivity and hysteresis. Our minimal model genes establish a regulatory circuit switching miR-155 expression, which in turn regulates the expression of inflammatory marker genes during inflammatory transitions.

## Introduction

Macrophages are essential innate immune cells, characterized by a high plasticity supporting their capacity to exert antagonistic functions in a time and stimulus-dependent manner. These cells transit between various inflammatory states in a tightly controlled process involving signaling as well as transcriptional and post-transcriptional regulatory networks. Identifying the molecules robustly controlling those dynamic transitions is crucial for elucidating the molecular basis of disease progression and for designing novel macrophage-based therapeutic strategies (Wang *et al*, 2014; Netea *et al*, 2016).

Identification of these molecules might be facilitated by the recognition that macrophage inflammatory transition is a bistable, or even multistable, system (Geiß *et al*, 2022). Transitions towards defined pro- or anti-inflammatory phenotypes have been shown to involve the same signaling pathway (Wang *et al*, 2014), indicating that their regulation is not linear. This suggests a system with the property of bistability or (more likely) multistability, referring to multiple steady state configurations defined by the concentrations of particular molecules. These steady states represent cellular attractors characterized by the self-sustained acquisition of a particular phenotype within the spectrum of macrophage activation states, which transitions are defined by systems-level properties. Accordingly, a macrophage in a particular steady state acquires an immunological memory (hysteresis) that affects the level of resistance to leaving this steady state (phenotype) and shapes the transitioning trajectories to other steady states(Murray *et al*, 2014). Examples of hysteresis in macrophages are found in trained immunity (Netea *et al*, 2016) or the lipopolysaccharide (LPS)-induced tolerance (Foster *et al*, 2007).

Studies of induced classical pro-inflammatory and alternative anti-inflammatory states indicate that pro-/anti-inflammatory transitions consist of bistable molecular switches of counteracting regulations (Wang *et al*, 2014; Murray *et al*, 2014; Biswas & Mantovani, 2010; Mills *et al*, 2000; Sica & Mantovani, 2012; Mathis & Shoelson, 2011; Ip *et al*, 2017). These molecular switches have been reported to emerge in several important regulatory processes of macrophage activation, dependent on various inflammatory mediators, signaling molecules, and transcription factors (TFs) (Palsson-Mcdermott *et al*, 2015). IRF/STAT signaling are central pathways in controlling macrophage inflammatory activation with the potential for the emergence of multistability. The pro-inflammatory regulators STAT1, STAT5, IRF5, SOCS3, NFKB-p50-p65, and HIF-1α have antagonistic and counteracting interactions with the anti-inflammatory regulators IRF4, SOCS1, STAT3, STAT6, NFKB-p50-p50, and HIF-2α. Several of these transcription factors, including NF-κB, AP-1, and STAT family members, also potentially participate in macrophage trained immunity, resulting in hundreds of times higher gene expression in a short window of time (Wang *et al*, 2014; Netea *et al*, 2016; Sica & Mantovani, 2012; Zhao *et al*, 2019). All these counteracting activities or inhibitory cross-talks between the pro/anti-inflammatory pathways indicate that inflammatory transitions may be considered as multistable systems consisting of molecular switches of mutually exclusive regulations. Further experimental and computational evidence of the emergent property of multistability in macrophage phenotypes has been reported, revealing the underlying principles of those molecular switches that include mutual-activation or mutual-repression circuits (Zhao *et al*, 2019; Berez *et al*, 2020; Nickaeen *et al*, 2019; Smith *et al*, 2016; Callard, 2007; Frank *et al*, 2021). However, the current models that describe this property are simplified and include less variables than realistic biological networks do, precluding a comprehensive analysis of the precise molecular mechanisms sustaining the switches that regulate phenotypic transitions.

Among these variables are microRNAs (miRNAs) which have not been widely included in these studies, despite being key regulators of gene expression. MiRNAs are crucial in modulating macrophage phenotypic transitions, as demonstrated by the tight link between miRNA deregulation and excessive or impaired inflammatory response (Essandoh *et al*, 2016). Moreover, they interact with transcription factors, which are key regulators and components of molecular switches. Transcription networks of miRNA-TF interactions have been reported to assemble complex network circuits and to target hubs leading to non-linear, systems-level properties such as bistability, ultrasensitivity, and oscillations (Ferrell & Xiong, 2001; Xiong & Ferrell, 2003).

We hypothesize that macrophage phenotypic transitions are controlled by molecular switches with the emergent property of bistability in miRNA/TF-regulated networks of gene expression. To prove our hypothesis, we used human monocyte-derived macrophages and followed a top-down systems biology approach facilitating the bioinformatic identification of the most relevant circuits controlling macrophage inflammatory transitions. Our approach uses experimental data (“top”) from system-wide analyses of mRNA, miRNA, and protein expression to reconstruct a large-scale biological network, followed by a parameterized mathematical model based on our biocomputational platform BioNetUCR 16 (“bottom”). Our multiomic data enabled the simplification of a large-scale network of interactions potentially regulating differentially expressed TFs and miRNAs in pro- and anti-inflammatory conditions. Due to the challenging task of parameter estimation for bistable models, we developed a novel parameter estimation pipeline. This new pipeline facilitates the construction of a bistable model of miRNA-transcription factor interactions able to undergo steady-state transitions triggered by transient perturbations in the abundance of specific genes or miRNAs. The analysis of the dependencies of these transitions led us to a minimal bistable model consisting of the interactions of miR-155 with four transcription factors, able to recapitulate the full model transitions and hysteresis. Finally, we confirmed our model experimentally using miR-155 mimics showing that increased levels of miR-155 trigger a pro-inflammatory transition, without changing expression levels of other model genes. With this approach, we contribute consolidating data for the recognition of macrophage inflammatory transitions as a bistable system and describe a potential role for a miR-155-mediated circuit in macrophage bistability. Altogether the data reported in this study contribute to the understanding of macrophage transitions for the future design of robust therapeutic strategies for cancer therapy or the treatment of inflammation-based diseases.

## Results

### Differential gene expression profiles of human macrophages under pro- and anti-inflammatory stimulations suggest the presence of properties related to bistability

Human macrophages treated with LPS+poly(I:C) or with IL-4+IL-10 have been previously reported to resemble a pro-inflammatory or an anti-inflammatory phenotype, respectively (Geiß *et al*, 2019). To characterize the kinetics of macrophage polarization in our model, we monitored a list of inflammatory marker genes by RT-qPCR including *IL1b*, *TNF, IDO1, CD163*, *MRC1*, and *SHPK*. Data were min-max normalized to balance the contribution of each gene and referenced to unstimulated macrophages (M0). Principal component analysis (PCA) was performed to reduce data dimensionality, and systems behavior was visualized plotting the first principal component (PCA1) as a function of time. We observed that pro-inflammatory stimulated macrophages (hereafter referred to as M1) reached their maximum polarization at 24 h whereas anti-inflammatory stimulated macrophages (hereafter referred to as M2) reached theirs at 72 h (Figure 1A, continuous red and blue thin lines, respectively), suggesting that both types of polarization have differential kinetics.

**Figure 1.**
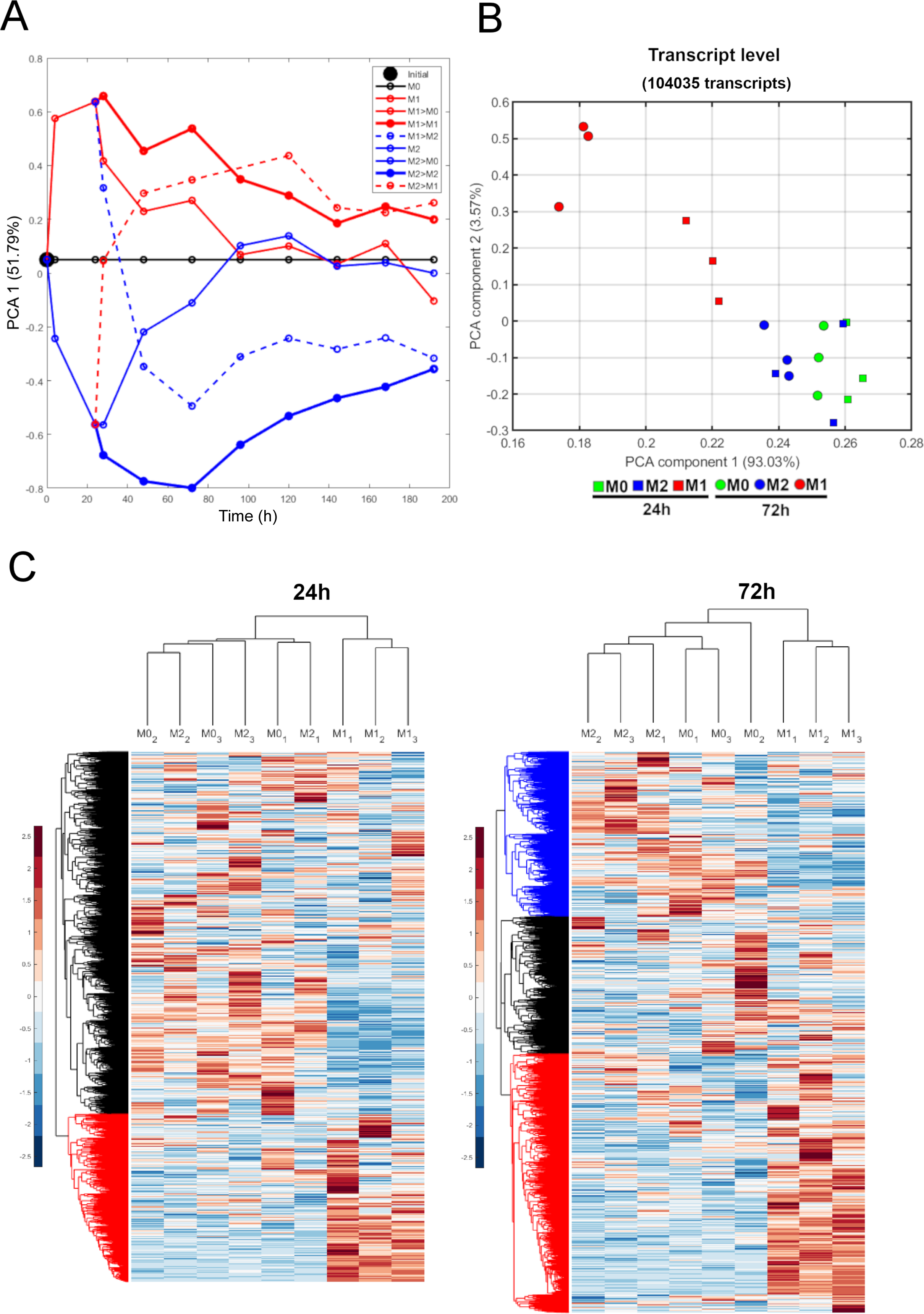
Differential gene expression of human macrophages under pro- and anti-inflammatory stimuli. **A.** Polarization and repolarization experiments of human macrophages treated with pro-inflammatory (M1, red) or anti-inflammatory (M2, blue) stimuli or left unstimulated (M0, black). The polarization experiment (first 24 h, continuous lines) is followed by stimuli removal (continuous lines), addition of the same stimuli (continuous bold lines) or a switch of the stimuli (dashed lines). The first principal component of the expression levels of inflammatory marker genes is shown as surrogate and as a function of time until 196. h. **B.** The two first principal components of transcriptomic expression data for the macrophages of three different donors at 24 h and 72 h after stimulation. **C.** Hierarchical clustering heatmap of gene expression of macrophages of three different donors at 24 h shows a clear cluster of M1 upregulated genes (red) and larger cluster of M2 and M0 downregulated genes (black). At 72 h, clear clusters of M1 upregulated (red), M2 upregulated (blue) and another cluster without a clear differential behavior (black) are identified.

To test whether macrophages retain any persistent memory of previous states able to impact their future repolarization to new states, we performed a repolarization challenge, where M1 macrophages were re-stimulated with the M2 stimuli (IL-4+IL-10) and the M2 macrophages with M1 stimuli (LPS+poly(I:C)) after 24 h (Figure 1A, dashed lines). As controls, we included a condition where the respective M1 and M2 stimuli were removed (Figure 1A, continuous lines) and another set of conditions where the cells were re-stimulated with the same original cocktails (Figure 1A, bold continuous lines). The data show that re-stimulated (repolarized) M1 and M2 macrophages do not reach a full transition to the maximal level of the opposite phenotype. The M2 macrophages restimulated with M1 stimuli (M2→M1) reached an M1-like phenotype close to that reached by M1 macrophages at 48 h after removal of the M1 stimuli (M1); it took another 48 h for these cells to reach an M1-level similar to that of M1 macrophages re-stimulated with M1 stimuli (M1->M1). Likewise, M1-macrophages re-stimulated with M2 stimuli reached their closest to M2-like phenotype at 72 h but this was still far from the maximum M2-phenotype (M2→M2). Although we cannot neglect completely the presence of leftovers of stimuli in the medium, this effect should be neglectable in these long-term experiments. Therefore, these observations are compatible with hysteresis, a property of bistable systems that retain a memory of previous states perturbing the behavior of future repolarizations.

To obtain a robust characterization of our experimental model we treated primary human monocyte-derived macrophages of three different donors for those maximum peaks of activation observed for M1 and M2 conditions (24 h and 72 h), keeping untreated macrophages as control (labeled as M0), and performed a full transcriptomic analysis as reported in materials and methods section. Principal component analysis of the full transcript data counts of 24 h showed that M1 macrophages cluster different to M0 and M2 macrophages showing dispersion along the y-axis (PC2) and a distinct separation along the x-axis (PC1; Figure 1B). In contrast, the M2 and M0 conditions are not completely separated, indicating that at 24 h those two conditions are rather similar, as expected from the culturing conditions of the M0 macrophages (Figure S1A, 24 h). For the 72 h time point, the cluster of the M1 macrophages is much tighter, suggesting the presence of a potential attractor of the M1 phenotype (Figure S1A, 72 h). In addition, the M2 and M0 macrophages show tighter clustering and could also be better distinguished by PC1, suggesting that they group around another potential attractor (M2/M0 phenotype). The presence of these attractors is more evident if a full PCA (including 24 h and 72 h samples) is plotted (Figure 1B). Here, it becomes clearer that the M1 samples at 24 h did not yet reach the M1 attractor observed at 72 h (upper-left corner), whereas the M2 and M0 samples remain closer to a potential M2 attractor (lower-right corner).

Next, we analyzed the transcript composition for both 24 h and 72 h and set our focus on protein-coding (most abundant) and long non-coding RNA (lncRNA)-coding transcripts (fourth in abundance) due to their functional importance (Figure S1). After separating the transcripts into those two classes, the PCAs for protein coding genes (Figure S1C) showed a similar behavior compared to the full transcriptome (compare Figure S2C with Figure S2A). However, at 72 h it was no longer possible to distinguish M2 from M0 condition at these first two components. These results suggest the existence of two different distinguishable attractors, one for the M1 and another one for the M2 and M0 conditions, becoming more evident at the later time point of 72 h. On the contrary, the PCA for the lncRNA transcripts showed a clear separation and clustering for the M0, M1, and M2 samples at 72 h, suggesting that differential lncRNA expression might mediate the differences between M0 and M2 conditions (Figure S1D). This underlines the role lncRNAs play in regulating immune cell functions (Walther & Schulte, 2020).

After collapsing transcript expression data into a gene level, we observed that the conditions were not well separated at 24 h (Figure S2A). However, at 72 h, the M1 samples could be better distinguished reinforcing the notion of an M1 attractor. This also confirms that M2/M0 conditions at both time points clustered around a second attractor, hereafter referred to as M2 attractor. These observations were further supported by our differential gene expression (DGE) analysis identifying 4004 differentially expressed (DE) genes between M1 and M0 conditions, and 3865 DE genes between M1 and M2 conditions after 24 h. In contrast, there are only 1112 DE genes between M2 and M0 conditions. Likewise, there are 3878 DE genes between M1 and M0 and 4400 between M1 and M2 with only 1237 DE genes between M2 and M0 conditions at 72 h. It is important to note that the log-fold change values in the annotation files (supplementary material) were calculated dividing M2 by M1 values (M1 is the reference), while in the comparisons of M1 and M2 they were calculated dividing M1 and M2 by M0 values (M0 is the reference).

The results of the hierarchical clustering heatmap for all genes further supported the observation of samples separation as well as the formation of a distinct expression pattern. (Figure 1C). Similar to what was observed at 24 h and 72 h using DGE analysis, there was a clearly distinguishable cluster for the M1-up regulated genes (red cluster) compared to M2 and M0. The opposite cluster of M2 and M0-upregulated and M1-downregulated genes could also be clearly observed at 72 h (blue cluster). At 24 h it is also possible to appreciate a large group of M1-downregulated genes but as expected it could not be further separated into M2 and M0 clusters (black cluster). Indeed, this hierarchical clustering analysis suggests that M1 macrophages have a very clear separation due to a robust group of differentially expressed genes whereas M2 and M0 macrophages are clustering around one attractor and cannot be distinguished based on our datasets (Figure 1C).

Given the observed behavior of the samples described above, we chose the time point of 72 h to perform a statistical functional over-representation test of the red (M1-upregulated) and the blue (M2/M0-upregulated) clusters of genes (Figure S2B). We included only GO biological processes with p-values lower than 0.05 with the Bonferroni correction for multiple testing. The first top 15 GO terms according to Fold Enrichment are shown in Figure S2B. Despite the differences encountered in M1 and M2 macrophages, there are some common GO-terms for both red and blue clusters such as cytoplasmic translation, cellular response to lipopolysaccharide, cellular response to molecules of bacterial origin, and cellular response to biotic stimulus, suggesting the presence of modules of co-regulated genes. The observed hysteresis between two potential attractors and these modules further supports the potential presence of bistability during macrophage polarization.

### Proteomics of M1/M2 stimulated macrophages point to mutually exclusive and time-dependent profiles of gene over-expression

Changes in the expression of protein coding transcripts led us to postulate the existence of potential attractors and hence of bistability. We therefore aimed to ascertain whether those changes in expression are also reflected at the protein level. Proteomic analysis of three macrophage preparations stimulated in a similar way as above resulted in triplicate forward and reverse experiment datasets. The log2 transformed data of the ratios between the different conditions are shown in Figure 2A. Although the identified genes presented in the diagonals are supported by the forward and reverse analysis, the identification of M1- and M2-regulated genes in this three-way comparison requires specific criteria as the fold-increases are larger in M1-than those of M2-upregulated genes and the M2 and M0 conditions present a similar behavior, as discussed above.

**Figure 2.**
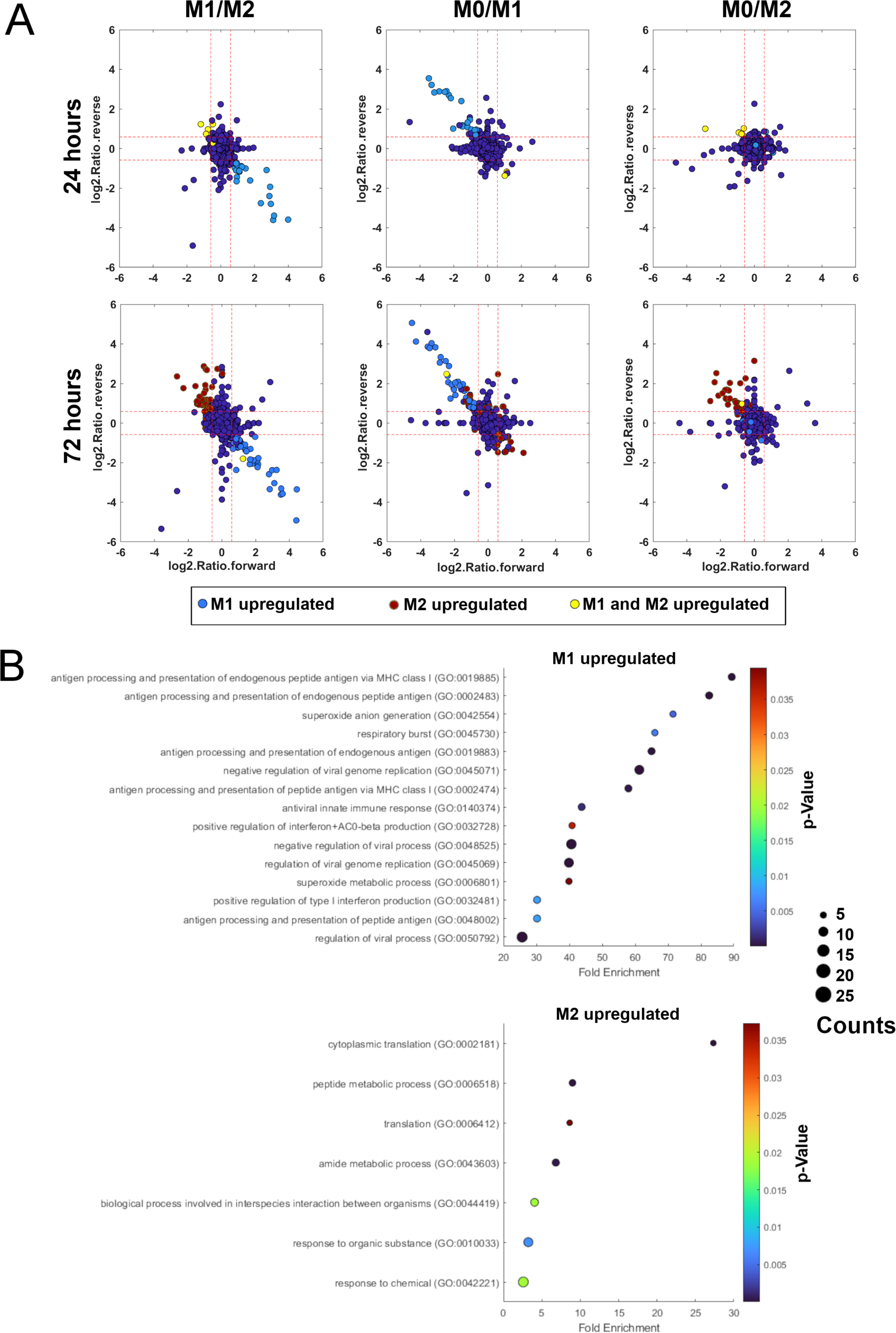
Differential gene expression profile at the proteomic level of stimulated macrophages. **A.** Log2 transformed data of the ratios between the different stimuli conditions for the forward and reverse experiments. After applying boolean operators, the identified M1-up regulated genes (light blue), M2-upregulated (brown) and M1+M2 upregulated (compared to M0, yellow) are shown. Downregulated genes are not labeled. **B.** Statistical functional overrepresentation tests of the M1 and M2 up-regulated genes at 72 h show different M1 and M2 programs at the protein level.

To improve the interpretation of these results and integrate the individual contribution of each experiment to a sensitive identification of M1 and M2-regulated proteins, we employed differential Boolean operators on the corresponding gene names across the different experiments. Since the M1 conditions lead to a very well separated cluster from M2/M0 at the transcriptomic level (see above) and they have higher fold-changes at the proteomic level, we used the ‘AND’ operator across the M1/M2 and M0/M1 experiments to increase the specificity of the identification of M1-up regulated genes (Figure 2A, light blue). In contrast, we used the ‘OR’ operator across the M1/M2 and M0/M2 experiments to increase the sensitivity of the identification of M2-upregulated genes (Figure 2A, brown) as they have lower fold changes and the differential expression between M2 and M0 is much lower compared to those to M1. The corresponding color labels of the M1 (blue) and M2 (brown) up-regulated genes are shown across all the experiments as well as those upregulated both in M1 and M2 (yellow) compared to M0 (Figure 2A). M1- and M2-downregulated genes were identified in a similar way by comparing to their corresponding M0 experiments (not labeled).

We could observe important differences between the conditions and time points. First, we observed time-resolved differences in M1 and M2 upregulation. For the M1 conditions there are 25 up-regulated genes at 24 h including the known M1 regulator STAT1 and other interesting M1 markers such as IDO1. The majority of those (24/25) are still up-regulated at 72 h together with 23 additional genes, suggesting that an early subset of M1-genes maintains its expression levels while activating additional ones at later time points, including other well-known M1 regulators such as STAT5A and STAT5B. On the other hand, there are only eight up-regulated genes in M2 conditions at 24 h and only four of them are still up-regulated at 72 h, but with 53 additional up-regulated genes at that later time point. Second, we observed the time-resolved differences in M1 and M2 gene down-regulation. In M1 conditions, we could observe only seven down-regulated genes, with three of them still down regulated at 72 h plus an additional 18 genes. Interestingly, six of those M1 down-regulated genes (compared to M0) correspond to up-regulated M2 genes at 24 h (*IGF2R, ITGA3, LPL, PSAT1, RPL13* and *RPL3*). Conversely, in M2 conditions there were only two down-regulated genes at 24 h and seven additional ones at 72 h. Of those, only the M2 downregulated *HLA-A* is also a M1-upregulated gene. The complete list of those target genes can be found in Supplementary Table 2. To further characterize the regulation of gene expression elicited by M1 and M2 conditions at the protein level, we performed a statistical functional overrepresentation test of the M1 and M2 up-regulated genes at 72 h, confirming that these differentially expressed proteins are related to different cellular functions (Figure 2B).

Altogether, these results show a time dependent increase in the number and strength of M1 and M2 up-regulated target genes but indeed only few down-regulated genes (compared to M0) suggesting that two alternative subsets of protein-coding genes are up-regulated for M1 and M2 conditions. Most importantly, all those genes corresponding to overexpressed proteins for both conditions are also differentially upregulated in our transcriptomic analysis. This observation is consistent with differential and mutually exclusive gene expression programs at the transcriptomic level for M1 vs M2 conditions, which are also reflected at the proteomic level.

### Multiomic data enable the simplification of a large-scale network of interactions potentially regulating differentially expressed transcription factors and miRNAs in M1 or M0/M2 conditions

The data reported above points to a possible bistability in gene expression that would characterize the transition of macrophages between different inflammatory states. We aimed to identify the role of miRNA-transcription factor interactions in the potential bistability of macrophage polarization by constructing a mathematical model to ascertain if bistability is theoretically possible in this system. For that purpose, we first constructed a network of regulatory interactions of the DE genes that potentially participate in the switches responsible for macrophage transitions. Our previously published biocomputational platform BioNetUCR enables the construction of such large-scale networks of gene-TF-miRNA interactions using inputs from different databases of experimentally validated regulators (Acón *et al*, 2021). We followed several steps for the construction and simplification of a potential network of miRNA-TF interactions involved in macrophage polarization:

#### 1. Definition of the list of transitioning genes of interest and their interactions

We first defined a list of genes, identified in our transcriptomics analyses, that are potentially switching between M1 and M2 attractors. To explore the full extent of gene expression changes across our experimental conditions, we plotted the logFC values of the differentially expressed genes in a unified plot where the changes in the ratios can be interpreted as changes in the expression of genes potentially switched (high to low or vice versa) across the conditions (Figure 3A). The genes without a statistically significant differential expression were assigned a zero for that specific transition. Here, we established a criterion to create a first list of M1-high/M2-low (M1 genes) genes including those having a positive logFC value in the M0 to M1 comparison and in the M2 to M1 comparison (green dots). We identified 1649 genes for 24 h (M1-24 h genes) and 1383 genes for 72 h (M1-72 h genes), with 1004 genes present at both times. We established a second criterion for the M1-low/M2-high (M2 genes) genes defined to include those genes with a negative logFC values for the M0 to M1 comparison and for the M2 to M1 comparison (yellow dots). We identified 1022 genes for 24 h (M2-24 h genes) and 1288 genes for 72 h (M2-72 h genes), with 610 genes being present at both time points. These criteria aim to identify genes transitioning from M0 and M2 to M1 regardless of their comparisons between M2 and M0, which are assumed to be in a very close or similar steady state as suggested by our previous results with the PCA analysis of the transcriptome (see Figures 1B, S1A and S2A).

**Figure 3.**
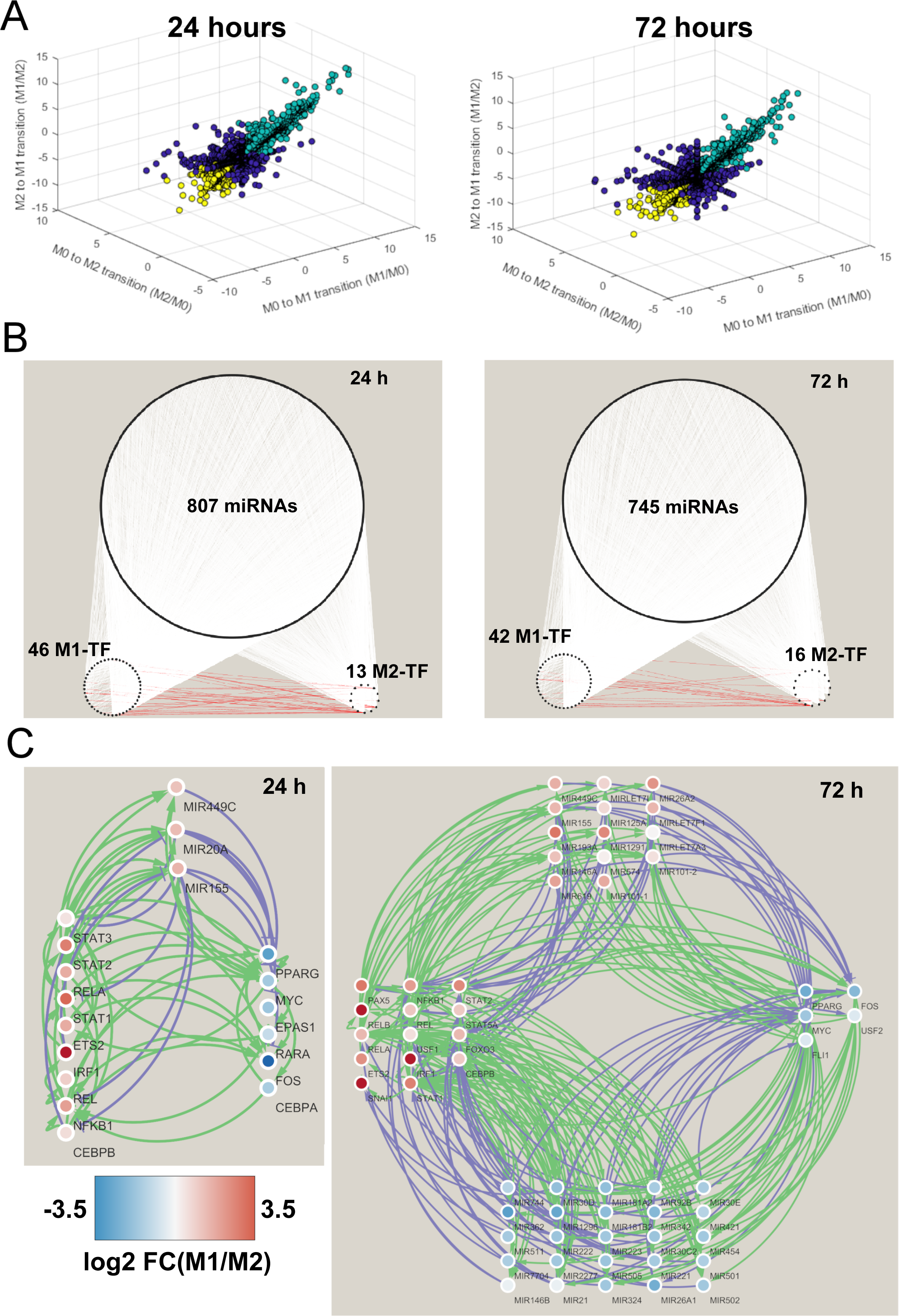
Construction and simplification of a network of transcription factor and miRNA regulators of differentially expressed genes based on multiomic data. **A.** Candidate switching-genes were identified based on hypothetical transitions between the different conditions and labeled based on logFC values of the differentially expressed genes. M1-high/M2-low (M1 genes) genes have a positive logFC value in the M0 to M1 comparison and in the M2 to M1 comparison (green dots). M1-low/M2-high (M2 genes) genes have a negative logFC values for the M0 to M1 comparison and for the M2 to M1 comparison (yellow dots). **B.** After several steps of construction and simplification (see text), large scale networks of miRNA and TF regulators were identified for the potential switching-genes. The network excludes those target genes unless they also have regulatory function (miRNA or TF) and the graphs indicate the direct interactions between the M1 and M2 transcription factors (red edges), as well as many potential interactions with miRNAs (white edges). **C.** Additional simplification steps based on proteomic and miRNA expression data led to identification of simpler networks where activating interactions (green) are counteracted by inhibiting interactions (purple) and complex circuits could be established between M1 (red scales of log2 FC(M1/M2) comparison) and M2 (blue scales of log2 FC(M1/M2) comparison) nodes.

#### 2. Initial network construction and simplification based on differential expression of transcription factors

Given the size of the potential network regulating gene expression in macrophage transitions, we first constructed a script in MATLAB (MathWorks) to look at our BioNetUCR database of experimentally validated interactions reported for transcription factors and miRNAs directly regulating the list of differentially expressed genes that are potentially switching between M1 and M2.Out of a total amount of 1835 miRs and 470 TFs available in the BioNetUCR database, we found 58527 interactions for the M1-24 h genes with other genes. This M1-24 h subnetwork includes 4075 nodes, 274 TFs, and 1838 miRNAs. 1559 genes (out of 1649) are potentially regulated either by TFs or miRNAs and 56 M1D1 genes also work as TFs. For the M2-24 h genes, we could map them to 29705 potential interactions with other nodes, where 946 (out of 1383) genes are targeted by miRNAs or TFs. Therefore, the M2-24 h subnetwork is constructed out of 4425 nodes including 188 TFs and 1806 miRNAs, and 23 M2-24 h genes also work as TFs. The union of these two subnetworks (M1 and M2) led to a full 24 h network of 87932 interactions. To reduce network complexity, we kept only those TFs that are also differentially expressed, reducing the list to only 56 differentially expressed M1-24 h-TFs and 23 M2-24 h-TFs. There is of course no intersection between the differentially expressed M1 and M2 TFs, suggesting that those TFs are involved in differential gene expression programs. In addition, we kept only miRNAs potentially involved in regulatory loops as they could be activated by at least one differentially expressed TF. Afterwards, we loaded the resulting network in BioNetUCR to remove sinks and sources left alone (degree of 1) to further reduce network complexity as they do not directly participate in regulatory circuit loops potentially accounting for bistability. Finally, we visualized this network in Cytoscape. As shown in Figure 3B, the network at 24 h has 866 nodes including 807 miRNAs and 59 differentially expressed TFs, 46 for M1 and 13 for M2 (Figure 3B, left). The graph shows direct interactions between the M1 and M2 transcription factors (red edges), as well as many potential interactions with miRNAs (white edges).

A similar approach was used for the construction of the regulatory network at 72 h. We found 47009 interactions for the M1-72 h genes with other genes. The M1-72 h subnetwork includes 3866 nodes, 191 TF and 1824 miRNAs. 1293 (out of 1383) are potentially regulated either by TF or miRNAs and 54 M1-72 h genes also work as TF. We could also map 37714 potential interactions for M2-72 h genes, where 1198 (out of 1288) genes are targeted by miRNAs or TF. The M2-72 h subnetwork is formed out of 4442 nodes including 181 TF and 1810 miRNAs, and 21 M2-72 h genes also work as TF. The union of the M1 and M2 subnetworks for 72 h led to a full 72 h network of 84723 interactions. Again, we reduced network complexity by keeping only the differentially expressed TF, narrowing down our list to 54 M1-72 h-TFs and 21 M2-72 h-TFs with no intersection between them. To simplify the network, we kept only those miRNAs activated by at least one differentially expressed TF as they could be involved in regulatory circuits. We loaded this network in BioNetUCR to remove sinks and sources not participating in regulatory circuits and generated the network for Cytoscape. The resulting network has 803 nodes including 745 miRNAs and 58 differentially expressed TFs, 42 for M1 and 16 for M2 (Figure 3B, right). The graph shows direct interactions between the M1 and M2 transcription factors (red edges) and many potential miRNA-interactions (white edges).

### 3. Network simplification based on differential miRNA expression

Since the complexity of these networks is still paramount, we ran a differential miRNA expression analysis for our experimental conditions to narrow down the list of potential miRNA regulators (see supplementary table 3). There are both upregulated and downregulated miRNAs for these macrophage transitions. At 24 h the comparison between M1 and M2 led only to 4 differentially expressed miRNAs: miR7704 is downregulated and miR449C, miR155, and miR20a are upregulated. miR155 is also differentially expressed when M1 is compared to M0. The comparison between M2 and M0 led to zero differentially expressed miRNAs. This list increases at 72 h as the comparison between M1 and M2 displays 18 upregulated and 23 downregulated miRNAs. A very similar list is observed by comparing M1 to M0 conditions obtaining 18 upregulated miRNAs and 28 downregulated miRNAs. For 72 h, the comparison between M2 and M0 led to 7 upregulated and 2 downregulated miRNAs. Noteworthy, the upregulation of miR155 and miR449C and the downregulation of miR7704 are observed already for 24 h and remain for 72 h, suggesting that those miRNAs could participate in early bistable switches potentially driving the switch of other subsequent circuits at 72 h. The higher number of differentially expressed miRNAs at 72 h suggest that those miRNAs may participate in the maintenance of those two potential steady states but the most important miRNA-regulated switches driving the transitions happen already at 24 h.

To select the most relevant miRNAs for both time points we included all miRNA differentially expressed across all comparisons (M1 vs M0, M1 vs M2, M2 vs M0). This led to a list of only four differentially expressed miRNAs at 24 h and 58 miRNAs at 72 h. After deleting sinks and sources (degree one), we obtained a 24 h network of 29 nodes and 97 edges, including three miRNAs and 25 TFs. The same procedure was done for 72 h leading to a network of 78 nodes and 401 edges, including 42 miRNAs and 36 TFs. 16/24 TFs of the 24 h network are also present in the 72 h network and miR155 and miR449C are also present in both simplified networks. These observations further indicate that some early circuits could be already active at 24 h and contribute to the maintenance of the steady states at 72 h.

#### 4. Further network simplification of transcription factors based on differential proteomics

Since many post-transcriptional modifications could take place until protein expression, we narrowed down the list of TFs to preserve only those with reported regulatory interactions with the list of differentially expressed genes identified both at the transcriptomic and proteomic level. This led to a reduced list of transcription factors, including 58 TFs at 24 h and 82 TFs at 72 h. Upon intersection of this list with that of the ongoing network simplified by miRNA interactions, we kept only those TFs robustly controlling our target genes at both gene and protein levels and able to interact with the differentially expressed miRNAs. For the 24 h time point, this led to a reduced network of 18 nodes and 66 edges including three miRNAs and 15 TFs (Figure 3C, left). For the 72 h time point, we obtained a reduced network of 58 nodes and 256 edges, including 39 miRNAs and 19 TFs (Figure 3C, right). Of these networks we plotted the log_2_ FC(M1/M2) comparisons to confirm M1 upregulation (red) and M2 upregulation (blue). It can be appreciated that many activating interactions (green) are counteracted by inhibiting interactions (purple) and complex circuits could be established between M1 (red scales) and M2 (blue scales) nodes with a potential switch-like behavior of bistability in the expression of modules of co-regulated genes transitioning at the same time from one potential steady state to another, driven by hub genes able to control the simultaneous transition of the whole module.

#### 5. Network simplification based on node involvement in regulatory loops

In our model, MYC and NFKB1 are among the most connected transcription factors and miR-155 is the most connected miRNA, participating in several types of putative regulatory motifs. Noteworthy are the positive feedback loops potentially leading to bistability between M1 and M2 TFs such as those formed by TF-TF interactions. For instance, that formed by MYC/ETS2 and other more complex motifs such as NFKB1>MYC>RARA>NFKB1, NFKB1>MYC>CEBPA>NFKB1 or STAT1>MYC>CEBPA>NFKB1>IRF1>STAT1. In addition, the presence of miRNAs leads to even more complex loops. However, more important than the number of connections of a node (degree within the network) is whether a particular node participates in potential regulatory motifs (complex positive/negative feedback loops) at least at the topological level (within the graph of interactions). We therefore implemented a script using the deep search algorithm in MATLAB (dfsearch) for the exploration of graphs corresponding to our networks. First, the stochiometric matrix was constructed for each network followed by the construction of the graph using digraph (MATLAB). Second, a deep search was performed starting from each node on those graphs and the ‘edgetodiscovered’ events were highlighted to find those edges ending a cycle (Figure S3 A and B). Within those discovered sinks we determined the shortest path (shortestpath, MATLAB) from each node to itself and filtered only for those interactions creating the circuit. After confirming the absence of replicated circuits, we filtered them to keep only those including both M1 and M2 nodes and determined the sign of their regulation (positive or negative) as the product of the signs of the individual regulations of each of the edges conforming the respective circuit. For the 24 h network, a total amount of 49 circuits coming back to their respective nodes were found, 41 of them including both M1 and M2 nodes. The level of involvement for each node in those circuits is shown in Figure S3C (24 h) including both negative feedback loops (blue) and positive feedback loops (red). For the 72 h network, a total of 228 circuits were identified, of which 224 involve both M1 and M2 nodes. The individual node involvement in those circuits is shown in Figure S3C (72 h).

This multiomic approach enabled network simplification for the discovery of two simplified networks of potential co-regulated genes under the control of complex regulatory circuits. The circuits identified for the 24 h network are also present within the 72 h network, suggesting that the most important transition-driving circuits are already functional at 24 h.

### A novel parameter estimation pipeline leads to the identification of a bistable model of miRNA-transcription factor interactions

Our results indicate that the identified networks contain modules of co-regulated genes switching from one state to another and under the control of circuits potentially leading to bistability. Indeed, many of these potential circuits include positive feedback loops at the topological level. However, their functionality as feedback-loops and their ability to enable bistability depends on the kinetic parameters of all their interactions. Therefore, the emergence of the systems property of bistability and the identification of robust targets controlling those potential regulatory circuits requires a quantitative systems biology approach to estimate the kinetic parameters of their interactions.

Because our gene expression data for M1- and M2-stimulated macrophages is limited to only two time points, we do not have information about the precise trajectories of the transitions between those two potential steady states. Nevertheless, we aimed to demonstrate whether it is theoretically possible to obtain one model of coregulated gene expression able to converge to at least two steady states, depending on the initial concentration of the species in the M0, M1 and M2 conditions. This model was based on the identified circuits within our 24 h network, as it most probably includes the earliest and most relevant targets to control macrophage transitions. Thus, we excluded 5 nodes (EPAS1, FOS, REL, RELA, STAT2) as they do not participate in any circuit (see Figure S3C) in our 24 h network, obtaining a further simplified network (Figure 4A) for model construction.

**Figure 4.**
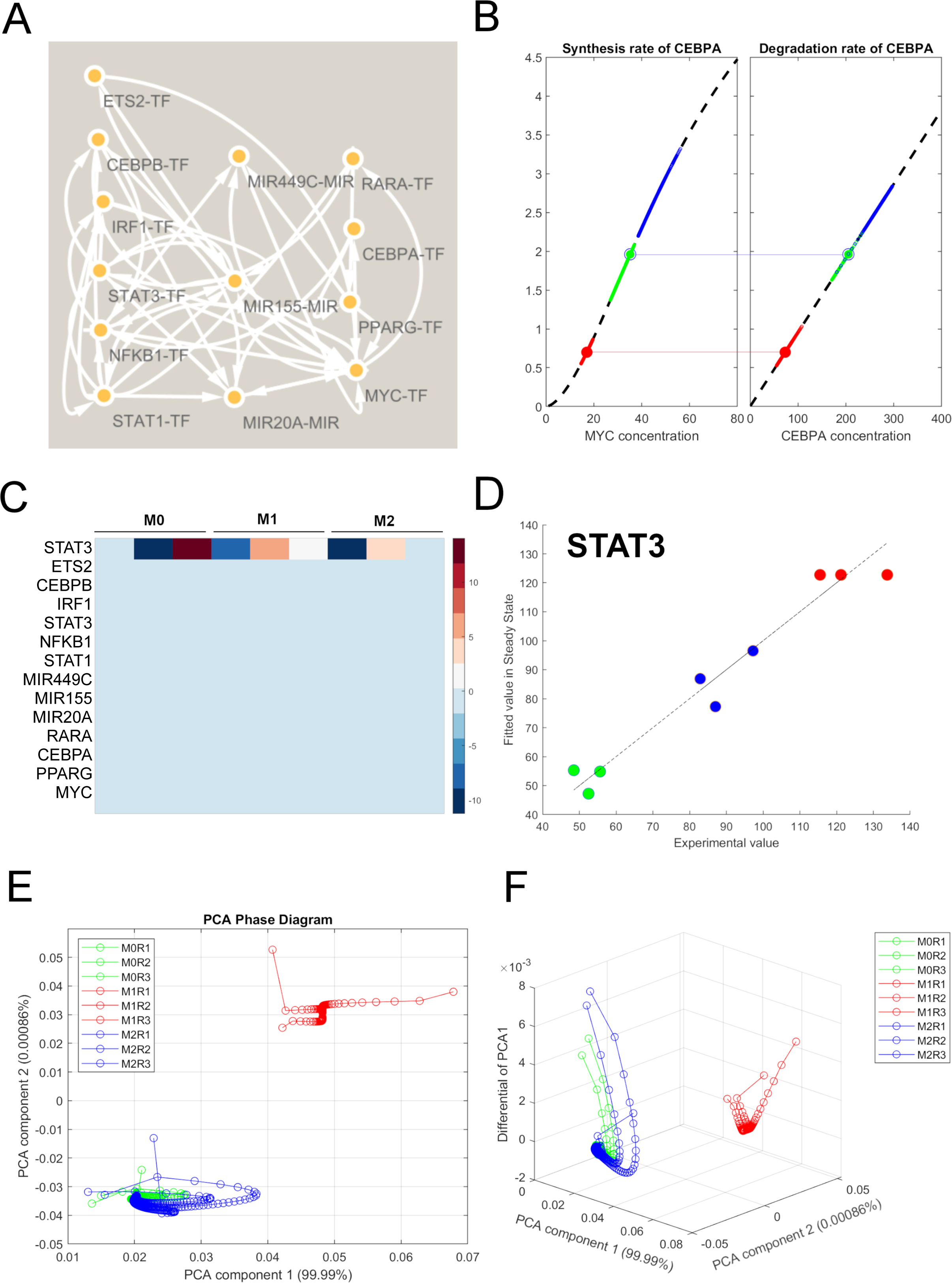
A bistable model for miRNA-transcription factor interactions was fitted using a novel parameter estimation pipeline. **A.** Simplified network version after removal of nodes that do not participate in any positive or negative regulatory loop. **B.** Graphical illustration of the fitting strategy for CEBPA. Synthesis depends on MYC concentration and degradation depends on CEBPA concentration. COPASI finds two steady state concentrations (large dots) where synthesis and degradation are equal. These concentrations are found within the ranges of the experimental data for the M0 (green), M1(red) and M2(blue) experimental conditions. **C.** Heatmap of the error (difference between experimental and simulated data) for the individual fitting of STAT3 as a function of its context. **D.** Direct fitted (simulated) and experimental data comparison for the individual fitting of STAT3 showing a good qualitative correlation and its compatibility with more than one steady state as a function of the remaining model species. **E.** The two first principal components of model species concentrations over time as surrogate to show that two different steady states of the model are reached depending on the initial experimental concentrations of M0 (green), M1(red) and M2(blue) macrophages. **F.** The two first components were plotted against the derivative of the first component to visualize the rate of change of the model asymptotically approaching the two different steady states. The intervals between two circles represent 1-time arbitrary unit.

BioNetUCR automatically constructs a mathematical model of differential ordinary equations from the network interactions including mass-action kinetic terms (and parameters) for synthesis, degradation, activation, and inhibition for each species. We included several modifications in BioNetUCR and in the COPASI model to ensure that we are able to model bistable systems (described in detail in Materials and Methods section). These modifications enabled us to increase the flexibility of our fitting strategy to determine if it is theoretically possible to obtain a bistable model for our two potential attractors based on the available circuits of the network.

Our experimental and simulated mRNA expression data are expressed in transcripts per million (TPM) as a function of gene copy number (two for all genes) and the measured mRNA concentrations of all the species (independent variables). To recapitulate the experimental data in our model, parameter estimation is required. However, the fitting of a bistable model is very challenging, hence we developed our own fitting strategy as a pipeline of systematic parameter estimation able to search for bistability. The general strategy is to find a set of parameters enabling synthesis and degradation rates of a particular species to intersect more than once. Therefore, we constructed a new model using a hill expression to obtain sigmoid behaviors of the synthesis reactions but kept linear degradation expressions for simplicity. This is also key to our fitting strategy since this degradation term of one species depends only on one degradation parameter (Kd) and on its own concentration.

The first step of our pipeline is to change the expression of the degradation constants (Kd) to an assignment of their respective synthesis flux rate divided by the experimentally expected concentration at steady state (in this case equal to the initial concentration for each experiment). Here, we ran parameter estimation with global and local algorithms correcting parameter boundaries until reaching an almost perfect fit (objective function close to zero). This is an artificially created model with multistability, as the model remains very close to the different initial concentrations when running simulations starting on the different initial concentrations of the individual experiments. Upon perfect fitting of this model, the Kd expressions were reverted to fixed values for parameter estimation either one by one (individual pipeline) or systematically to achieve a fully fitted model (full model pipeline).

First, we applied our pipeline as an individual fitting strategy for each of the species. We illustrate hereby the method with CEBPA as its synthesis rate only depends on MYC concentration and its degradation only depends on CEBPA itself. Thus, we can plot simple graphs of CEBPA synthesis and degradation rates as a function of only one species. After the reversion of the Kd expression, we repeated the fitting process until no boundary alerts were found and a low objective function was reached. We could confirm the presence of a ‘bistable-like’ behavior after running steady-state simulations starting from the initial concentrations of the different experimental conditions (M0, M1, M2) and verifying that the model indeed reached different final concentrations. Using these values for the equation’s parameters, we plotted the resulting CEBPA synthesis and degradation as a function of their MYC and CEBPA concentrations respectively, as well as the trajectories of time-course simulations starting from the different M0 (green), M1 (red), and M2 (blue) conditions, and the positions of the two identified steady states (large dots) (Figure 4B). As observed, the simulations with the starting M1 conditions reached a steady state where synthesis and degradation intersect at low MYC and CEBPA concentrations, very well within the range of the three M1 experiments (red dot on red trajectories). On the other hand, all simulations starting at M0 and M2 conditions (six experiments) reached a single steady-state where synthesis and degradation intersect at high MYC and CEBPA concentrations (green/blue dot near green/blue trajectories).

Subsequently, we applied this individual fitting strategy to isolate each individual species and to determine if a bistable behavior could be compatible with the surrounding network context and concentrations of all remaining species. These results were visualized for STAT3 by plotting the error for each of the experiments upon reversion of the Kd expression (Figure 4C). As illustrated, the remaining species have very low error, so this method enabled us to isolate STAT3 behavior by fixing the rest of the model species to the concentrations of the experimental data. The error of STAT3 was minimized with parameter estimation and compared to the experimental data in Figure 4D. The analysis of the data distribution indicates a good qualitative agreement between simulated and experimental data but also suggests that the concentration range of STAT3-regulators may enable more than one steady state for STAT3. The process was repeated for all individual species, where in general we could observe a good agreement between simulations and experimental data except for ETS2 and MIR449C (Figure S4). For most of the remaining species at least two possible steady states were identified, one corresponding to M1 conditions and at least one for M2/M0 bconditions. These results suggest that most species in the current network are compatible with bistable-like behavior enabled by its surrounding network species, except for ETS2 and MIR449C. This fitting strategy enabled us to discard those species, which potential bistability cannot be explained by the network, further reducing model complexity to include only those able to support each other’s bistable behavior.

After removing ETS2 and MIR449C from the network, the reconstructed model was used for what we have called “progressive full model fitting” (see materials and methods). Briefly, we first reverted the expressions of the Kds for fixed values to be estimated one by one, starting with those presenting the highest degradation fluxes, and repeating parameter estimation each time. Once a model species was well fitted, we verified the bistable behavior by running steady-state and time-course simulations. If a bistable behavior was identified, next we removed all the corresponding parameters of that species from the parameter estimation task to prevent further estimations from altering the behavior of the already fitted/bistable species. This process was systematically followed until all the Kds were fitted and determined. Finally, a fine tuning of the model fitness was performed by re-estimating the list of parameters of individual species contributing the most to the objective value, one by one, until a fully fitted model was obtained. To finally test the bistability of the whole model we ran steady-state and time-course simulations confirming the presence of two different steady-states for all species. To visualize this behavior, we plotted the two first principal components of the species concentrations out of time-course simulations, starting at the initial concentrations of the different experimental conditions (Figure 4E) and also plotted the absolute value of the first derivative of the PCA1 to visualize the rate of change of the system for homogeneous time steps (the intervals between circles) (Figure 4F). The results show that the simulations starting in the M1 conditions (red) converge into one steady state, whereas all the simulations starting in M2 and M0 conditions (blue and green) converge into a second steady state.

Our results indicate that we have developed a novel parameter estimation pipeline to successfully fit bistable models with potential applications to other systems. Moreover, our model indicates that it is theoretically possible to obtain a bistable behavior with our network of circuits involving the switched M1 and M2 coregulated genes. This also suggests that the two attractors observed in our previous data are indeed two different steady states of the system and they are reached depending on the current concentration of the interacting molecules. Therefore, such a system would potentially enable all these co-regulated miRNAs and transcription factors (and their targets) to simultaneously switch in the presence of external stimuli exerting an effect only on some key molecules, which can drive macrophage phenotypic transitions.

### Steady-state transitions of the bistable model can be triggered by transient perturbations of transition-driving genes but depend on the activity of other transition-enabling genes

Thus far, we have confirmed that it is theoretically possible to fit a mathematical model with the property of bistability after removing non-compatible species (in our case ETS2 and MIR449C) and applying a novel pipeline of systematic parameter estimation. The resulting network has 11 nodes (Figure 5A) and its model can reach two different steady states depending on the given initial concentrations of the experimental conditions (M0, M1, M2). To investigate whether our bistable model can switch between steady states upon transient perturbations of individual concentrations, we performed parameter scans of the initial concentrations for each individual species (see material and methods).

**Figure 5.**
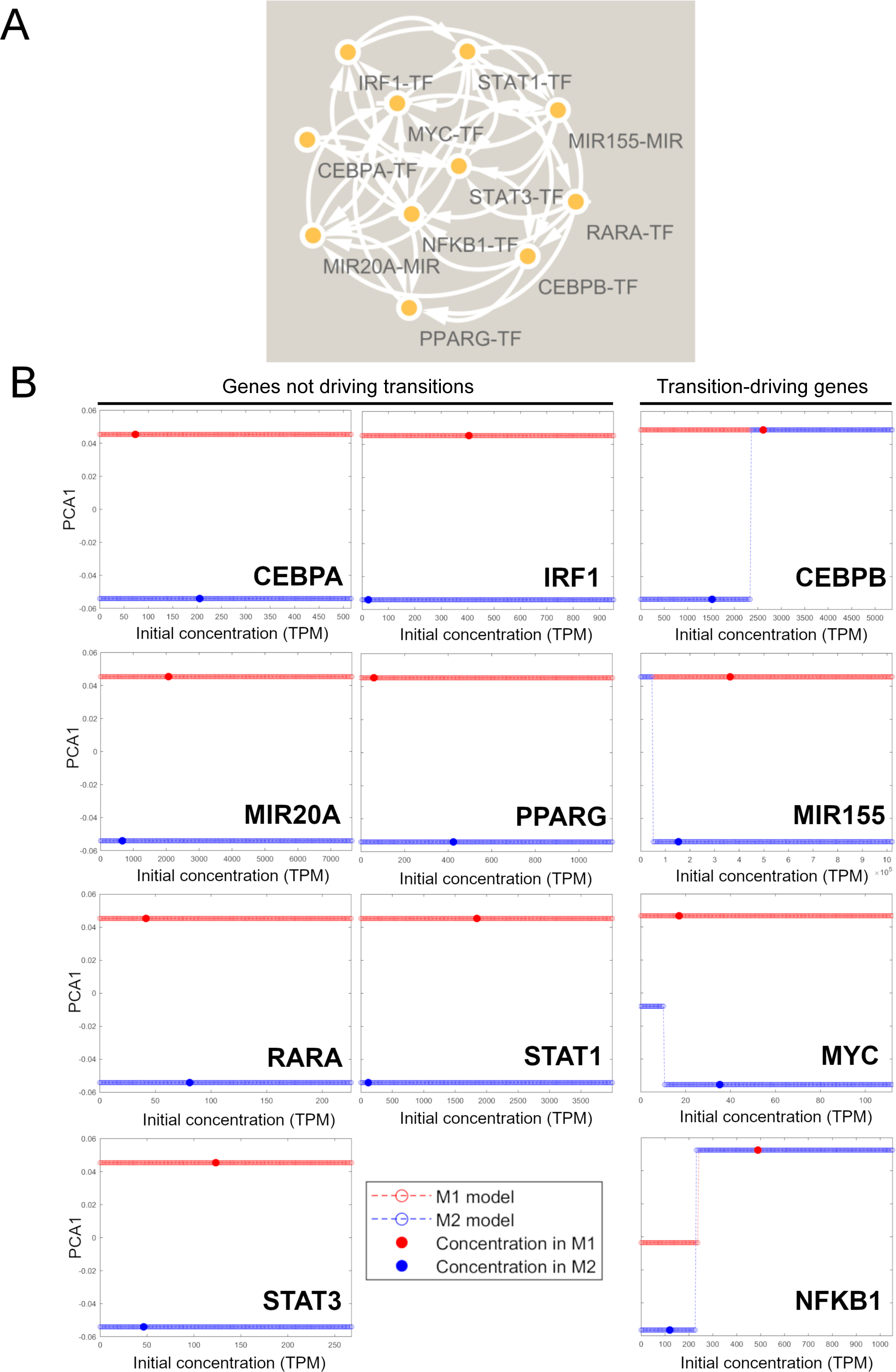
The bistable model of miRNA-transcription factor interactions obtained after progressive fitting can transition between steady states upon transient supercritical perturbations in the concentrations of several species. **A.** Further simplification of the fitted model after removal of the non-compatible species with bistable behavior. **B.** First principal component as surrogate of all the model species concentrations to monitor system status upon parameter scan of the initial concentration of individual species. The parameter scan was performed on the M1 model (red) and the M2 model (blue) to visualize model transitions between steady states. The species concentration at the original M1 and M2 steady states are depicted as large solid dots.

To perform the parameter scans (subtask steady-state), we started with the model in M1 (red) and M2 (blue) conditions by setting the initial concentrations to the corresponding steady-states. For both models, we scanned the effect of varying the initial concentration of each species from 0 up to two-times the maximal value reported in our experimental data and indicated the initial species concentration with a larger dot for the corresponding M1 or M2 steady state (Figure 5B). This parameter scan simulates a transient perturbation of the system where only this initial concentration is changed, and the model will then evolve to the closest attractor until reaching again a steady state. We plotted the first principal component of the model species concentrations as a surrogate to monitor the system status for the parameter scans of each species. The transient perturbation of the initial concentrations of CEBPA, IRF1, MIR20A, PPARG, RARA, STAT1, and STAT3 led the model to reach the same original steady state, registering no transitions. However, model transitions between the two steady states were registered for the remaining species. A transient increase in CEBPB in the M2 model led to a transition from M2 to M1 but the opposite behavior is not observed when CEBPB concentration is decreased in the M1 model. Similarly, the increase of NFKB1 in the M2 model triggers a transition from M2 to M1 whereas the decrease of NFKB1 in the M1 model triggers another transition from M1 to a third intermediate steady state. These two NFKB1-driven transitions occur at similar but not identical NFKB1 concentrations.

Please note that these CEBPB and NFKB1-driven transitions were observed within the concentration ranges of the experimental data and within the concentrations range defined between the M1 and M2 steady states. Nevertheless, outside those ranges two other transition trajectories emerged: for MIR155 a further decrease in the levels of MIR155 in the M2 model triggers a transition from M2 to M1, whereas no effects are noticeable for perturbations of MIR155 concentrations in the M1 model. Likewise, the decrease of MYC concentration in the M2 model (beyond the levels of the M1 steady state) leads to the transition from M2 to that same third steady state observed for NFKB1 (defined by the final concentrations of the species), whereas no transitions are observed for the perturbation of MYC in the M1 model (Figure 5B). This asymmetrical behavior in M1-to-M2 and M2-to-M1 transitions supports the concept of hysteresis, suggesting that the corresponding trajectories are different for both types of transitions.

Moreover, the response of the system to single perturbations indicates that transients in the concentration or activity of single genes could trigger the switch of the system from one steady state to another. However, it is questionable whether all those genes are required for the model to transition from one steady state to another or if there is only a minimal circuit of genes required. To answer this question, we performed additional in silico experiments using our models. We repeated the identified model transitions and evaluated the role of single species by fixating their initial concentrations of the corresponding models, which renders those species insensitive to changes. The parameter scans under the different perturbations are shown and the obtained trajectories were compared to the control conditions (Figure S5). The NFKB1-driven M1-to-M2 transition depends on the changes in CEBPB, IRF1, MIR155, and STAT1. On the other hand, the NFKB1-driven M2-to-M1 transition depends on IRF1, MIR155, and STAT1. The CEBPB-driven M2-to-M1 transition depends on NFKB1 and MIR155, whereas the MIR155-driven M2-to-M1 transition depends on IRF1, NFKB1, and STAT1. Finally, the MYC-driven transition depends on all the other species as any perturbed species leads to a blockade in that transition. Since this latter steady-state was not represented in our experimental data, we focused our study on the M1-to-M2 and M2-to-M1 transitions. In this regard, our results indicate that a circuit composed of the transition-driving species (NFKB1, CEBPB, MIR155) in addition to two transition-enabling genes (STAT1 and IRF1) can capture most of the behavior of this model. Indeed, if those transition-driving genes represent robust targets to induce switches in the steady states, the remaining transition-enabling genes represent interesting targets to potentially lock the system in a determined steady state.

### A minimal bistable model of macrophage M1/M2 transitions depending on the interactions of MIR155, NFKB1, IRF1, STAT1, and CEBPB presents hysteresis

To confirm that a minimal model of macrophage transitions could be established with those transition-driving and transition-enabling genes, we constructed and fitted a new model including only CEBPB, NFKB1, IRF1, STAT1, and MIR155 (Figure 6A). We followed the same procedure for parameter estimation as described in the previous section. The resulting model was able to recapitulate a bistable behavior as observed after setting the initial simulation conditions to the values of the different replicates (R) of macrophage treatments for both 24 h and 72 h (Figure 6B). The PCA representation of the model trajectories for each experiment shows that all M1 conditions reached a single steady state regardless of the time of treatment (red trajectories). In addition, all M0 and M2 conditions also reached a single different steady state independently of the incubation time (blue and green trajectories).

**Figure 6.**
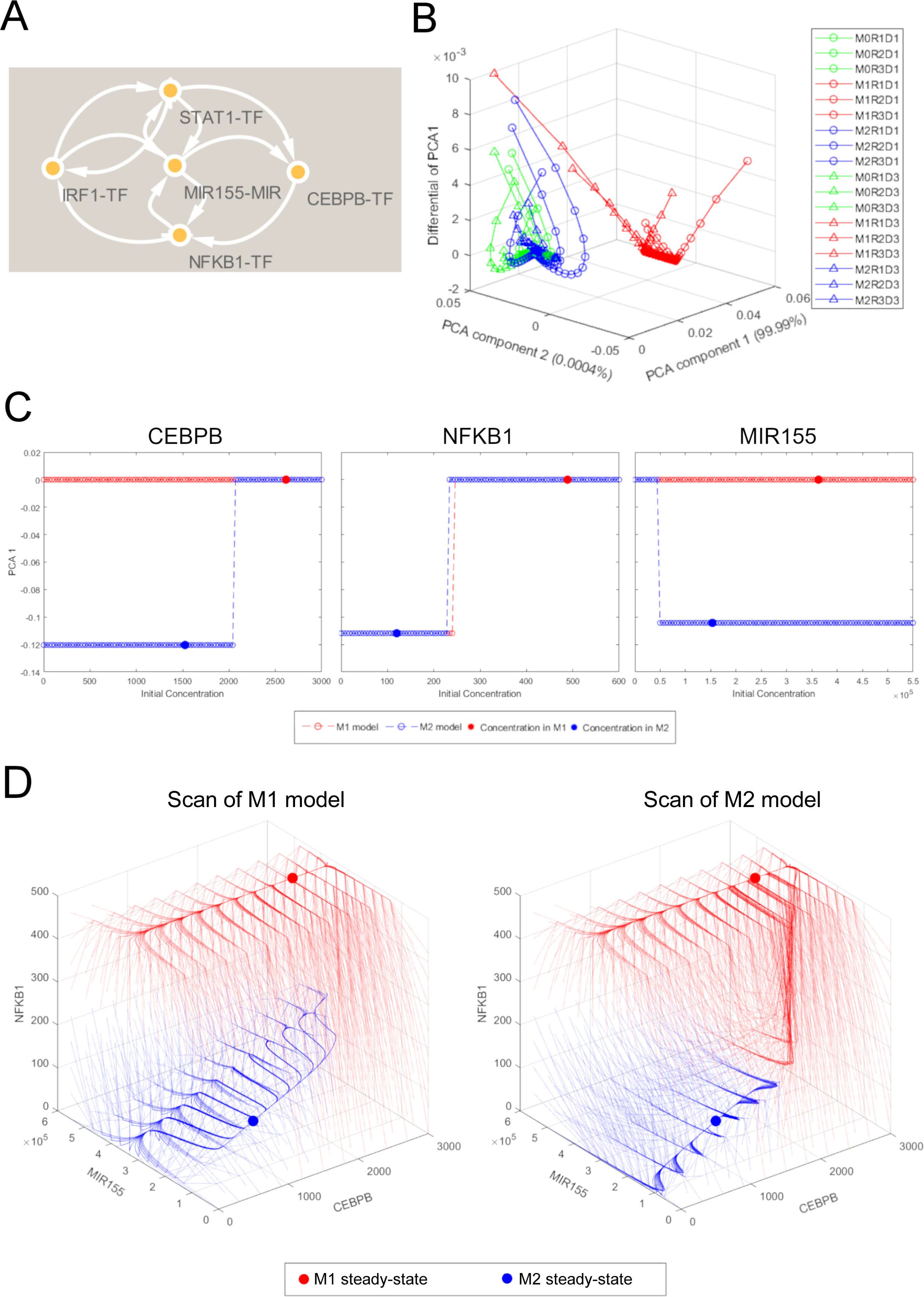
Hysteresis is observed in a minimal model of macrophage M1/M2 transitions. **A.** Final simplification of the minimal model after removal of non-required species for model transition between steady states. **B.** After parameter estimation of the minimal model, it was evaluated at the initial concentrations corresponding to the experimental data observed for 24 h and 72 h, It could be shown that all M1 samples (red) converged to a single steady state, whereas all M0 and M2 samples converged into another. The PCA1 and PCA2 are used as surrogates for system status and the differential of PCA1 is included to show the rate of change of the system towards the steady state. The interval between circles represents 1 arbitrary time unit. **C.** The minimal model recapitulates the four transitions driven by transient perturbations in the initial concentrations of the transition-driven species. The first principal component (PCA1) is shown as surrogate of system status as a function of a parameter scan of the initial concentration of the corresponding species. Large dots represent steady concentrations for the M1 (red) and the M2 (blue) models. **D.** Trajectory landscape of time course simulations as a function of the initial concentrations of NFKB1, MIR155 and CEBPB in a three-dimensional parameter scan of the M1 and the M2 model. The final steady state concentrations are shown as large dots, and the trajectories are colored according to their outcome.

Moreover, we aimed to evaluate whether this minimal model could also recapitulate the transitions observed between steady states upon perturbations of the concentrations of single species. For the CEBPB-driven transition, we could detect a M2-to-M1 transition upon the increase in CEBPB, but no M1-to-M2 transition upon a decrease in CEBPB. Likewise, both NFKB1-driven transitions were observed for the perturbations of the minimal model as for the full model. Furthermore, the MIR155-driven M2-to-M1 transition was also observed upon further decrease in MIR155 levels way below the M2 conditions, probably by releasing its inhibition on NFKB1 and CEBPB, and triggering thereby a switch to M1 steady state, in consistency with the full model (Figure 6C). To further illustrate the minimal model behavior during these transitions, we ran a time course simulation of the model in steady state and applied events with transient perturbations of the species concentrations. In all cases the first event took place at 300 time-units of simulation time by setting the species concentration to a subcritical change (below that required for transition), to illustrate how the system comes back to the same steady state. The second event of transient perturbation took place at 4000 time-units with a supercritical concentration change of the species (beyond that required for transition) to illustrate how the system switches to a different steady state upon transition (Figure S6A). These results indicate that our minimal model is bistable and can recapitulate the observed transitions of the full model between the M1 and M2 steady states.

We also ascertained the dependency of these model transitions on other species by fixing their concentrations to the current steady state and repeating the parameter scans to observe if the transitions are abolished, confirming thereby that the model cannot be further reduced. As expected, the CEBPB-driven M2-to-M1 transition and the NFKB1-driven M1-to-M2 transition require all the remaining species to be unfixed. Nevertheless, the MIR155-driven M2-to-M1 transition and the NFKB1-driven M2-to-M1 transition occur despite the fixation of CEBPB concentration to low M2 values (Figure 6SB). This difference between M1-to-M2 and M2-to-M1 transitions suggests that those transitions undergo different trajectories, and therefore a potential hysteresis could emerge in this system. Moreover, for this model to recapitulate all the observed potential transitions, it cannot be further reduced. This result confirms that this is indeed the minimal model required to explain the molecular switches for the steady state concentrations of these species.

Since only three species can drive the transitions between steady states in this minimal model, we could now plot the response to a tri-dimensional parameter scan of single and combined transient perturbations to their concentrations. The resulting trajectories of the model in response to perturbation drew a sample of the full landscape of transition trajectories (Figure 6D). This landscape shows the distribution of critical threshold concentrations able to trigger the switch from one steady state to another, starting either from M1 conditions (M1 model) or from M2 conditions (M2 model). Interestingly, a clear difference can be observed for several trajectories as a few of the same perturbations switched the M1 model to M2 but also switched the M2 model to M1. These differential trajectories were filtered and plotted in Figure 6SC, suggesting that the perturbation landscape has a region of hysteresis, where the final steady state depends on the previous history of the system.

### The experimental expression profile of the minimal model genes recapitulates the behavior of inflammatory marker gene expression in macrophage phenotypic transitions

To ascertain whether our theoretical full and minimal models of macrophage transitions could explain part of the biology of the macrophage experimental transitions, we repeated the first repolarization experiment until 72 h and performed PCA (Figure 7A). The first principal component of the assessed M1/M2 inflammatory marker genes and of the genes involved in our full model were plotted over time (Figure 7A left and right, respectively). The changes in inflammatory marker expression reproduce the macrophage polarization starting in M0 (black) into M1 (red) and M2 (blue) states until 24 h. As expected, upon re-stimulation at 24 h the polarized macrophages started trajectories in the opposite direction of their original phenotypes. The M2-to-M1 repolarized macrophages reached an M1-like phenotype already after 4 hours of repolarization, whereas the M1-to-M2 repolarized macrophages did not reach a complete transition even at 72 h, suggesting some degree of hysteresis from the M1 polarization. After confirming this repolarization behavior, we monitored from the same samples the gene expression of nine genes coding for the species of our full model. Although the M2-to-M1 repolarization was slower, those reprogrammed M2 macrophages reached a full M1 phenotype after 72 h. In contrast, the M1-to-M2 repolarized macrophages did not reach a full M2 phenotype at 72 h, showing a similar behavior to the expression profile of the marker inflammatory genes (Figure 7A).

**Figure 7.**
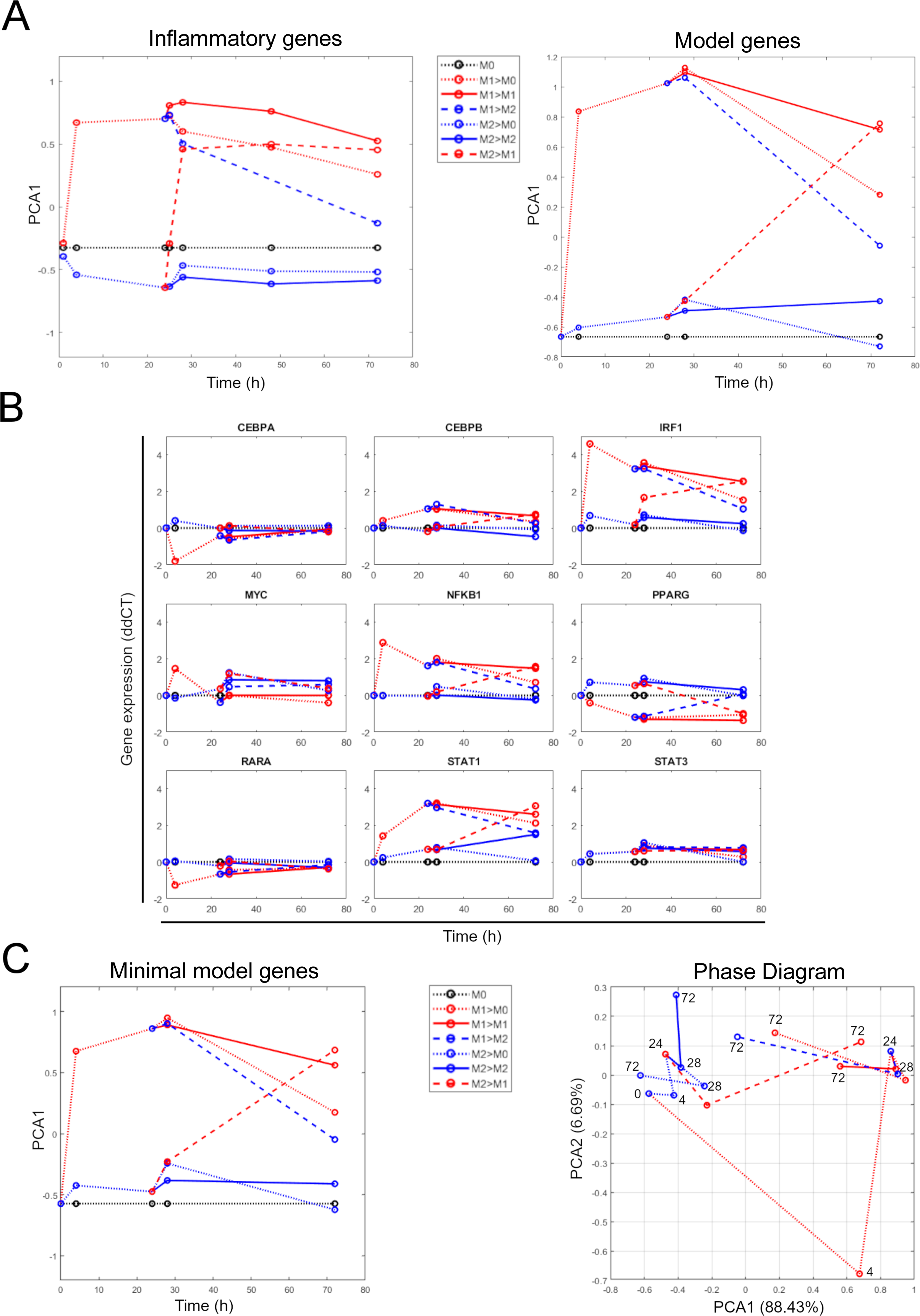
The experimental behavior of marker gene expression is reflected by the changes in gene expression of the minimal model species. **A.** A comparison of the overall gene expression profile between inflammatory marker gene expression and the experimental expression of the full model species in a polarization-repolarization experiment of macrophages. Data is represented as the first principal component of gene expressions as surrogate of macrophage status over time. **B.** Gene expression profiles for all the genes of the full model over time for the same experiment. The minimal model species present the most similar behavior to the overall PCA profile for both the model species and the inflammatory marker gene expression. **C.** A principal component analysis was done for the expression profiles of only the minimal model species to confirm that its profile resembles that of the inflammatory marker gene expression over time (left). The first two components are shown (right) to better illustrate the distances between the different conditions across time.

To dissect this behavior at the single gene level, we plotted the changes in expression for all those single genes and noticed that those that are part of the minimal model (CEBPB, IRF1, NFKB1, STAT1) reflect the overall gene expression behavior including the same repolarization pattern (Figure 7B). Indeed, when we repeated the PCA analysis with only those four genes we obtained a profile very similar to the overall behavior of the system (Figure 7C left). A plot of the two first components better illustrate the distance changes in expression for these four genes, starting at 0 h and taking the different trajectories depending on the stimulation conditions (Figure 7C right). These results suggest that this set of four genes can reproduce the behavior of inflammatory marker gene expression during macrophage repolarization experiments.

### An increase of miR-155 perturbs the repolarization of pro-inflammatory macrophages

Based on our current model we aimed to evaluate the role of miR-155 perturbations on macrophage transitions. Therefore, we simulated the transition of a macrophage from M0/M2 to M1 and back to M0/M2 by changing the concentrations of NF-KB beyond the critical values for transition (NF-KB driven transition). The control condition shows the expected behavior where the expression of all genes transitioned from a low to high concentration as the systems moves towards the M1-steady state (the concentration of NF-KB is set at 200 after 1700 s of simulation). In addition, the concentration is set back to 200 at the simulation time of 5000 s, leading to a decrease of their concentrations back to the M0/M2 concentrations as the system returns to that steady state (Figure 8A). This scenario simulates in silico our experimental control condition where a macrophage is set to M1 polarization and then to M2 repolarization. As shown in Figure 6C, an increase in miR-155 concentration does not induce a M2-to-M1 transition, neither does the decrease in miR-155 induce a M1-to-M2 transition.

**Figure 8.**
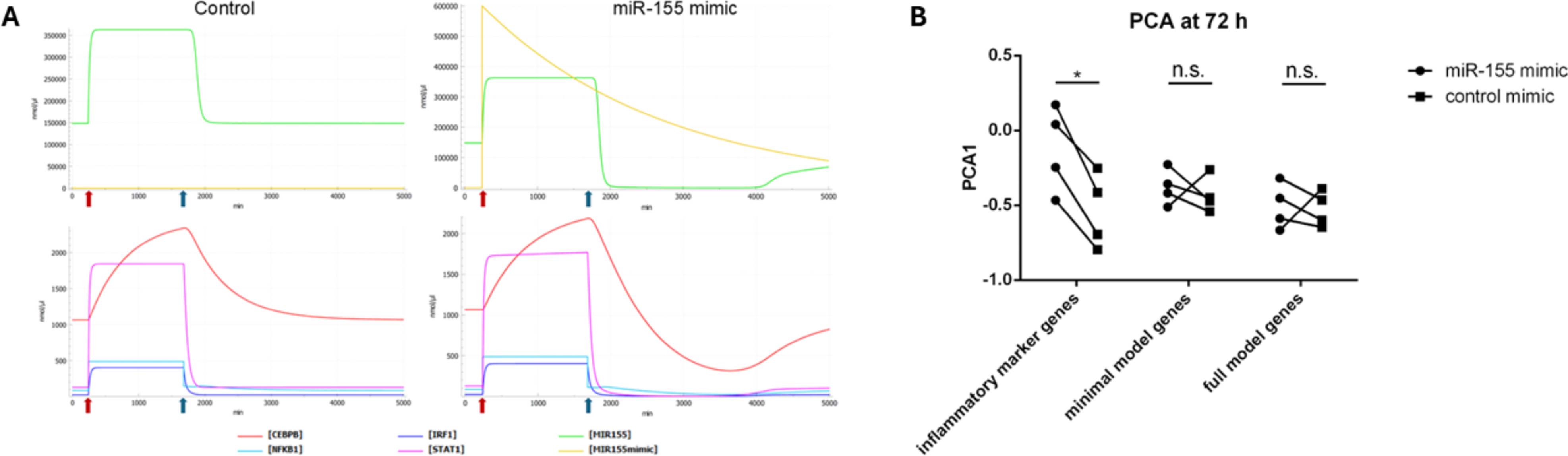
Addition of miR-155 mimic is not shifting concentrations of model genes, but of inflammatory genes. **A.** Modeling of miR-155, CEBPB, NFKB1, IRF1, and STAT1 expression levels after pro-inflammatory stimulation (red arrow) followed by anti-inflammatory re-stimulation (blue arrow) depending on the addition of control mimic (left) or miR-155 mimic (right). **B.** First principal component of inflammatory marker genes, minimal model genes, and full model genes at 72 h after addition of miR-155 mimic or control mimic. A higher PCA1 value reflects a more pro-inflammatory state. T-test using the Holm-Sidak method, with alpha = 5% was applied. * = significant difference with p = 0.006.

But we wanted to know if a miR-155 perturbation could influence the scenario of NF-KB-driven transitions depicted above. Therefore, we induced miR-155 perturbations in silico to assess its role in these transitions, by simulating the addition of an artificial miR-155 (mimic) and plotting the in silico abundance of genes based on our minimal model (Figure 8A). Since miRNA mimics are chemically modified to increase stability and reduce degradation, we used a low Kd (degradation constant) for the miR-155 mimic. Of note: to determine the start concentration and validate a very slow degradation of the mimic after transfection, we checked their abundance via qPCR 0 h and 72 h after transfection. As shown in Figure 8A after pro-inflammatory stimulation (red arrow) concentrations of all minimal model genes and miR-155 increased reaching the M1-steady state but decreased again after anti-inflammatory repolarization reaching the M2-steady state (blue arrow). The addition of miR-155 mimic (Figure 8A right) led to decreased levels of endogenous miR-155 and lowered levels of the minimal model genes after anti-inflammatory stimulation. However, this effect is transient as the model genes finally stabilized at the steady state concentrations, indicating that the transitions were completed.

To validate our model hypothesis, we transfected macrophages with miR-155 mimic or the corresponding control and repeated the repolarization experiments, focusing on M1-to-M2 polarization based on the results shown in Figure 8A. We performed PCA (Supplementary Figure S7) and plotted the first component of the inflammatory marker genes, the minimal model genes, and the full model genes at 72 h time point (Figure 8B). Going in line with our mathematical model, we observed only minimal changes looking at the model genes, but a clear pro-inflammatory trend looking at the inflammatory marker genes. The higher PCA1 value of the macrophages treated with the mimic (dots) compared to their controls (squares) indicates a more pro-inflammatory profile of these cells. This finding indicates that the minimal model genes establish a module that controls miR-155 concentration and is able to sense external stimuli (i.e. through the activation of NF-kB) and drives macrophage transitions. This module is robust to a transient increase of miR-155, however the concentration of miR-155 is a critical factor (i.e. through the regulation of downstream targets) defining the transient macrophage phenotype as indicated by the changes in inflammatory markers.

## Discussion

Bistability, the ability of a system to rest stably in one of two distinct states, is a fascinating phenomenon that offers both robustness and sensitivity. Indeed, biological systems often operate in noisy environments where fluctuations in molecular concentrations or external signals are common. Bistable systems are inherently robust to such noise, as the system tends to remain in one of the stable states unless a strong enough perturbation occurs. Bistability also enables switch-like, ultrasensitive response to critical signals (Ferrell & Ha, 2014; Bagci *et al*, 2006). This is the case for gene regulatory networks, where a gene is turned "on" or "off" in response to a triggering signal. Bistability will ensure that the resulting state is maintained even after the initial triggering signal is removed. This kind of memory is essential in processes like immune response (Kadelka *et al*, 2018).

Despite its importance, detecting bistability in biological systems remains challenging due to the complexity and dynamic nature of regulatory networks, inherent biological noise, and experimental limitations. First, these systems often involve multiple, overlapping feedback loops that can obscure bistable dynamics (Alon, 2007). Second, transient states or graded responses may mimic bistability, requiring long-term or perturbation-based assays to confirm true bistable behavior (Ferrell & Xiong, 2001). Third, stochastic noise and cell-to-cell heterogeneity can blur state distinctions, complicating the interpretation of population-level data (Eldar & Elowitz, 2010; Raj & van Oudenaarden, 2008). Moreover, accurately detecting bistability often demands perturbations and precise measurements that may be experimentally unfeasible (Ozbudak *et al*, 2002). Bistable behavior can also depend on narrow biochemical conditions, if these are missed, bistability may go undetected (Ferrell & Machleder, 1998). Finally, interpreting such behavior requires computational models capable of capturing nonlinear dynamics and feedbacks inherent to biological regulation (Tyson *et al*, 2003).

To investigate bistability in human macrophage polarization, we first analyzed the expression kinetics of established inflammatory markers via RT-qPCR. In polarization–repolarization assays, we observed bimodal gene expression distributions compared to unstimulated macrophages (M0). Notably, pro- and anti-inflammatory phenotypes remained stable for up to 192 h in the presence of stimuli. Upon repolarization, we observed a hysteresis-like effect, particularly in the M1-to-M2 transition, where gene expression did not fully revert to M2 levels even by the experiment’s end. While residual cytokine effects cannot be completely ruled out, the return to M0 conditions 72 hours post-stimulus suggests minimal carryover.

These results support the presence of bistability in our model and suggests the existence of hysteresis in macrophage polarization. The latter, however, remains debated. Some studies in mouse macrophages report full reversal to M0 within 96 hours and no hysteresis, while others argue that macrophages may respond flexibly to immediate environmental changes, minimizing hysteresis effects. Conversely, several reports support hysteresis, citing asymmetric repolarization dynamics, epigenetic memory, and the influence of prior stimuli on future responsiveness. We propose that this debate is partly due to a lack of mechanistic insight into the regulatory circuits underpinning bistability—an area our study addresses through minimal circuit modeling.

To further explore bistability, we performed transcriptomic analysis on human monocyte-derived macrophages, capturing gene expression at peak M1 and M2 activation times (24 h and 72 h). Different donors represent biological replicates to consider the inter-individual noise in gene expression. Despite the limited number of samples, we observed tight clustering of M1 samples at 72 h, suggesting a potential M1 attractor, while M2 and M0 samples clustered around a second attractor, suggesting that M0 cells are already engaged in a M2-like state. These two potential attractors could be observed at the levels of full-transcripts, protein-coding transcripts, and gene expression. Hierarchical clustering revealed two distinct gene expression patterns: one cluster of M1 upregulated genes (Figure 1C, red cluster) and another cluster of M1 downregulated genes (Figure 1C, blue cluster), indicating potential co-regulation. Functional over-representation analysis confirmed this hypothesis and showed that some genes in both clusters were involved in similar processes, albeit regulated differently in M1 versus M2 or M0 states. Proteomic analysis, however, revealed a distinct program for M1- and M2-upregulated genes, with many transcriptomic changes not reflected at the protein level Therefore, we used a multiomic approach for network construction and simplification. We identified transcription factors (TFs) potentially controlling M1 and M2 gene expression at 24 h and 72 h. By filtering for differentially expressed TFs with proteomic targets, we found distinct regulatory patterns for M1 (46 TFs at 24 h, 42 TFs at 72 h) and M2 (13 TFs at 24 h, 16 TFs at 72 h), further supporting the idea of separate gene expression programs.

These findings, along with the evidence of hysteresis and potential co-regulated modules, suggest bistable behavior in macrophage polarization. To investigate this further, we built a mathematical model of gene expression regulation using a top-down systems biology approach, focusing on the role of miRNA-mediated regulatory circuits. With this in silico approach, we identified miRNA-mediated circuits, focusing on miRNAs like miR-155 (upregulated in M1 at 24 h) and miR-146a (upregulated in M2 at 72 h), known to play a role in macrophage transitions. This regulatory network, including complex circuits formed by transcription factors and miRNAs, further supports the presence of bistability in our macrophage model.

To confirm the presence of regulatory circuits in these networks, we ran a deep search algorithm to identify positive and negative feedback circuits and to quantify the degree of involvement of each species within these circuits, observing that most nodes participate in more than one of such circuits. Our multiomic approach simplified the network, revealing two key modules of co-regulated genes controlled by complex feedback circuits with the potential for bistability. Notably, these circuits, identified at 24 h, were also present at 72 h, suggesting that the initial circuits are functional at 24 h and contribute to the amplification of the transition across the entire macrophage gene expression profile. These early circuits likely enable bistability, with the transition being largely committed by 24 h. A quantitative characterization of the initial circuits at 24 h could lead to the identification of such initial bistable circuit robustly controlling macrophage transitions.

We developed a novel parameter estimation pipeline consisting of a series of steps for an easier fitting of potentially bistable models. We used this pipeline to explore whether a model of our identified network could theoretically enable bistability, under the assumption that all the network nodes undergo transitions between our hypothetical M1 and M2 steady states, defined by the fitting to the experimental data. The first step of this pipeline is to achieve the maximum level of simplification at the level of the network topology. Therefore, we chose to start the model construction for the 24 h network after removing those nodes that do not participate in any positive or negative feedback loop, as detected by our deep-search algorithm. The second step is to construct the model with Hill expressions enabling a sigmoid-like behavior of the synthesis functions while keeping the degradation as a linear mass action kinetics function. This will enable an easier search of the parameters enabling that the synthesis and degradation function intersect more than once. The third step is to change the expression of the degradation constants (Kd) to an assignment of their respective synthesis flux rate divided by the experimentally expected concentration at steady state and run the parameter estimation until an almost perfect fitting to the experimental data is achieved. Under these conditions, only the synthesis function of each species is sensitive to its regulatory context (experimental concentrations of regulating species) assuming that they all undergo a transition between those two steady states. The fourth step is to revert the Kd expression to fixed values for the parameters to be estimated either individually (individual pipeline) or systematically (full model pipeline).

However, this process is not straightforward since returning each Kd to a fixed value reduces the flexibility as now the degradation and the synthesis functions must intersect at only one value for each steady state, instead of intersecting at each experimental value. Therefore, we recommend to start fitting those species with the highest degradation fluxes and using a combination of global and local algorithms to find the best fit in each case. In addition, it was important to verify that the fitted species conserved a bistable behavior running steady-state and time course simulations after each parameter estimation. It was also important to remove the parameters of the fitted species before running the parameter estimation for the next species. Otherwise, the fitting and the bistable behavior of the previous species was lost. In this way, we could partially overcome the limitation of the high-dimensional parameter spaces, since we split the parameter space for each of the model species as a function of its context (i.e. regulating species concentrations), obtaining thereby a good approximation of the model to a single solution despite having multiple data points. We consider that this approach also contributes to parameter identifiability as it is very easy to lose the correct fitting if the corresponding species parameters are not removed from the next parameter estimation task. Moreover, after this pipeline is systematically followed to revert all the Kds and estimate all the corresponding species-related parameters, it is still possible to perform a fine tuning of the parameter estimation. Since now a single steady-state concentration is obtained for each of the species regulating others, we performed such fine tuning by re-estimating the list of parameters of individual species, starting by those presenting the highest error. Therefore, we have developed a novel parameter estimation pipeline to successfully fit bistable models with potential applications to other systems overcoming limitations such as the high parameter space, the non-linearity of the system, the noise in biological data and the computational complexity. We consider that this approach could also be adapted for parallel computing for the modeling of large-scale networks as the optimization problem can be divided for each species. Moreover, this platform will enable the identification of bistable models that can be further refined against time course data, including the measurement of the full transition trajectories to obtain precise time-resolved bistable models able to describe the system’s transitions.

Using this approach, we constructed a mathematical model able to present the property of bistability, where two different states were identified depending on the initial conditions of the model. The model identified transition-driving genes, such as CEBPB, NFKB1, and miR-155, that exhibit ultrasensitivity to concentration changes, triggering transitions between steady states. This ultrasensitivity suggests potential hysteresis, as the system’s response to perturbations depends on its previous state. Additionally, STAT1 and IRF1 were identified as transition-enabling genes that facilitate the transitions driven by the transition-driving genes.

We constructed a minimal model with only those five genes (NFKB1, miR-155, STAT1, IRF1, and CEBPB) that could explain macrophage polarization transitions. This simplified model confirmed bistability, as it converged to distinct attractors for M1 versus M0 or M2 conditions. The model also exhibited hysteresis, as transitions depended on the prior state. Experimental validation supported the model, as the gene expression profiles of those genes mirrored inflammatory marker gene expression in polarization-repolarization experiments. Additionally, the addition of artificial miR-155 sustained a more pro-inflammatory state but did not prevent the M1-to-M2 transition. Our model explains this behavior: the minimal model gene circuit has the emergent property of bistability, defining thereby two alternative steady states and therefore two alternative extremes in the concentrations of miR-155 in response to external stimuli through the activation of NF-kB through signaling (NF-kB-driven transitions). Transient perturbations in miR-155 are insufficient to switch the whole circuit to an alternative steady-state, but the activity of the miR-155 on other downstream targets alters the expression of other inflammatory markers to a certain extent. These findings indicate that the function of the bistable circuit is indeed to provide a stable behavior for the concentration of miR-155 (and the other genes), but this concentration is the critical factor that will finally influence the macrophage phenotype.

In conclusion, we present evidence for bistability during macrophage phenotypic transitions. The direct interactions of our minimal model genes assemble a circuit where several transition-driving genes (NFKB1, CEBPB and miR-155) and transition-enabling genes (STAT1 and IRF1) can simultaneously switch from one steady state to another. The transition-driving genes could respond to external perturbations, able to change their transient concentrations or activities, such as signal transduction. If this perturbation surpasses a critical concentration, it will trigger the transition from one steady state to the other involving the whole circuit. Several reports provide robust evidence for the role of NF-κB in bistability, demonstrating how feedback loops and other regulatory mechanisms within the NF-κB signaling pathway contribute to the system’s ability to exhibit bistable behavior, which is essential for cellular decision-making processes (Hoffmann *et al*, 2002; Shinohara *et al*, 2014). On the other hand, also miR-155 has been implicated in bistable circuits (Chen *et al*, 2013) and was also shown to regulate inflammatory cytokine production in tumor-associated macrophages by targeting CEBPB (He *et al*, 2009). STAT1 multistability has also been implicated in macrophage polarization due to its direct interactions with STAT6 (Frank *et al*, 2021). In addition, IRF1 has been shown to govern the interferon-stimulated gene responses in human monocytes and macrophages as it is required to regulate chromatin accessibility (Song *et al*, 2021), and it is essential to STAT1-dependent immunity to mycobacteria (Rosain *et al*, 2023). Nevertheless, only a few studies report a potential role of miR-155 in NF-KB bistability in macrophages.

Interestingly, there is probably a temporal asymmetry between miR-155 and miR-146a expression during macrophage activation (Figure S8), creating a combined positive and negative feedback circuit controlling NF-KB activity, enabling a robust but time-limited inflammatory response (Mann *et al*, 2017). However, those reports of individual genes potentially present the property of bistability with very relevant functions but there was no report on how they could be integrated into a single circuit. Our results, therefore, expand the current knowledge on the role of this regulatory circuit on the bistability of miR-155 and those other potentially co-regulated genes, which are very important for the proinflammatory responses of the M1 macrophages. Our full model also offered a potential explanation for their coregulated downregulation in M2 macrophages and the coregulated upregulation of other important genes for the M2 phenotype, such as MYC, CEBPA, RARA, STAT3, and PPARG.

This model has been simplified in several ways to be able to fit in and identify potential bistable circuits, which clearly limits the identification of all the potential circuits actively engaged in the M1 and M2 gene expression programs. As shown in Figure 3, the potential networks and the number of circuits increase significantly at 72 h. Within this network, several other miRNA-mediated circuits could be present and have important regulatory roles for other macrophage subtypes, not included in our experimental model. However, it is more realistic to assume that macrophage activation represents a spectrum of different phenotypes, which are usually depicted as a continuum (Mosser & Edwards, 2008). Nevertheless, we hypothesize that due to potential multistability, this spectrum is probably not a continuum, but it is built up by many discrete steady states with overlapping phenotypes that could be identified at the molecular level as differential network attractors, defined as self-maintained steady states and controlled by many different bistable circuits. Consequently, it is very challenging to identify robust therapeutic targets able to control those transitions among the spectrum of inflammatory states without a full understanding of these multiple attractors.

Future work using single cell data will enable the identification of more potential attractors within the spectrum of macrophage activation and most importantly the trajectories across those steady states. The identification of trajectory-driving genes and the generation of pseudotime data sets will be instrumental in the construction of a full multistable model of macrophage activation. Despite the limitations of our approach, our work offers an initial proof-of-principle about the role of miRNAs in macrophage polarization, widens the repertoire of tools for the fitting of complex bistable models and offers the potential for the design of therapeutic strategies to trigger macrophages activation in cancer or deactivate them in proinflammatory diseases.

## Methods

### Monocyte isolation and differentiation into macrophages

Buffy coats were purchased from the Transfusion Center of the University Medical Center of the Johannes Gutenberg University (Mainz, Germany) and were obtained from anonymized healthy blood donors. All buffy coats used in this study are residual biological materials made available by the Transfusion Center to scientists on a randomized basis. Blood samples were collected and processed in accordance with the relevant German guidelines and regulations. Personal data are neither collected nor shared for this material. Peripheral blood mononuclear cells were isolated from buffy coats using a Ficoll gradient. Monocytes were differentiated into macrophages by culturing them for one week in cDMEM [DMEM (Gibco), 10% heat-inactivated FCS (Sigma-Aldrich), 2 mM L-Glutamine (ThermoFisher), 10 μg/ml Gentamicin (Sigma-Aldrich)] supplemented with 20 ng/ml macrophage-colony stimulating factor (BioLegend) under standard conditions (37°C, 5% CO_2_). Monocyte-derived macrophages were further cultured in the absence of M-CSF for another week prior to the experiment.

### Cell culture experiments using human macrophages

Human monocyte-derived macrophages were seeded in multi-well plates (Sarstedt) in cDMEM and cultivated overnight at 37°C and 5% CO_2_. Before stimulation or transfection at the indicated time points, medium was removed, cells were washed two-times with PBS and fresh cDMEM was added to cells. Cells were stimulated with established inflammatory stimuli (Geiß *et al*, 2019), either with the pro-inflammatory M1 cocktail [10 µg/ml polyinosinic-polycytidylic acid (poly-I:C, InvivoGen) + 10 ng/ml lipopolysaccharide (LPS, Sigma Aldrich)] or the anti-inflammatory cocktail [10 ng/ml IL-4 + 10 ng/ml IL-10 (BioLegend)]. For miR-155 manipulation experiments, cells were transfected with miR-155 mimic (mirVana, Invitrogen) or control mimic using Lipofectamine 2000 (Invitrogen).

### Total RNA isolation, cDNA transcription, and RT-qPCR gene expression profiling

After incubation, cells were washed with PBS, and total RNA isolation was performed using ReliaPrep™ miRNA Cell and Tissue Miniprep System (Promega) according to the manufacturer’s instructions. RNA concentration and quality were determined using a Nanodrop 2000c (ThermoFisher). For mRNA to cDNA transcription the FastGene Scriptase II - Ready Mix (NIPPON Genetics Europe) was used. Quantitative PCR (qPCR) analysis was performed using commercial probe-based primers (PrimeTime™ Gene Expression Assays, Integrated DNA Technologies) and a CFX Opus Real-Time PCR System (Bio-Rad Laboratories). Briefly, for each well of the 96-well qPCR plate (Sarstedt), 10 μl of qPCRBIO Probe Mix (NIPPON Genetics Europe) were mixed with 10 ng cDNA, 1 μl of the appropriate primer (Table 1), and filled up to 20 µl with H_2_O. Relative quantification (RQ) of gene expression was determined using the 2^−ΔΔCt^ method (Livak & Schmittgen, 2001). *SDHA* and *GAPDH* were used as reference genes. For miRNA transcription, 10x M-MuLV-buffer (New England Biolabs [NEB]), 1 mM adenosine triphosphate (Life Technologies), 1 mM dNTP-Mix (NEB), 10 U M-MuLV reverse transcriptase (NEB), 0.1 U poly(A) polymerase (NEB), 5 pM cel-mir-39 as spike-in control and 0.5 µM of reverse transcriptase primers (Table 2) were added to 25 ng total RNA in a total volume of 20 µl. The reverse transcription protocol consisted of 30 min at 37°C, followed by 60 min at 42°C and a final incubation for 5 min at 65°C. 5 µl of iTaq Universal SYBR Green Supermix (Bio-Rad Laboratories) were mixed with 0.56 ng cDNA and 0.5 mM of the appropriate primers (Table 2). SNORD110 was used to normalize miRNA results. Correctness of amplicons was verified using melt-curve analysis or agarose gel electrophoresis.

**Table 1:**
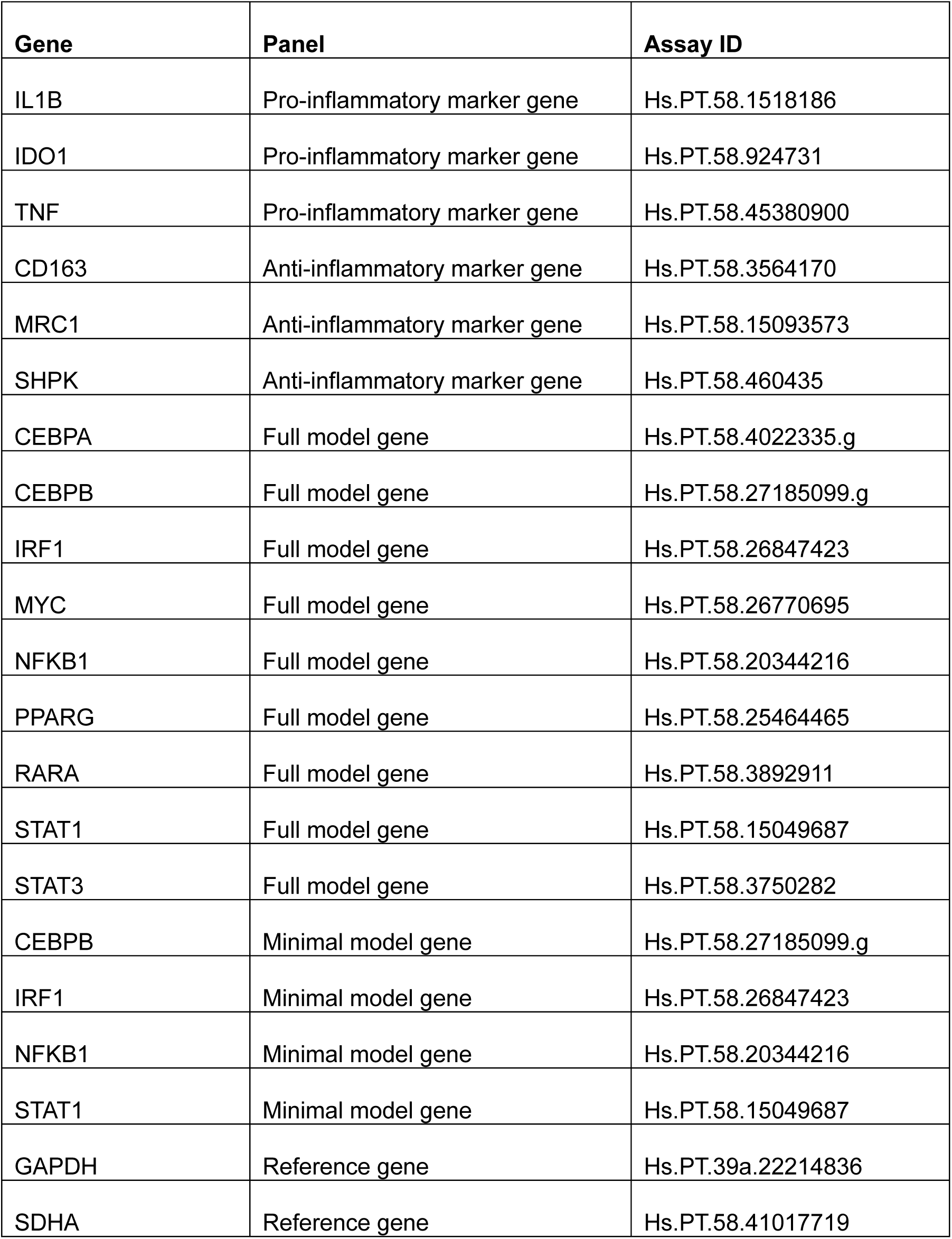
Genes (HGNC Symbols) used for RT-qPCR analysis, their corresponding panel, and assay ID (Integrated DNA Technologies). Genes defined either as pro- or anti-inflammatory are labeled as M1 or M2, respectively.

**Table 2:**
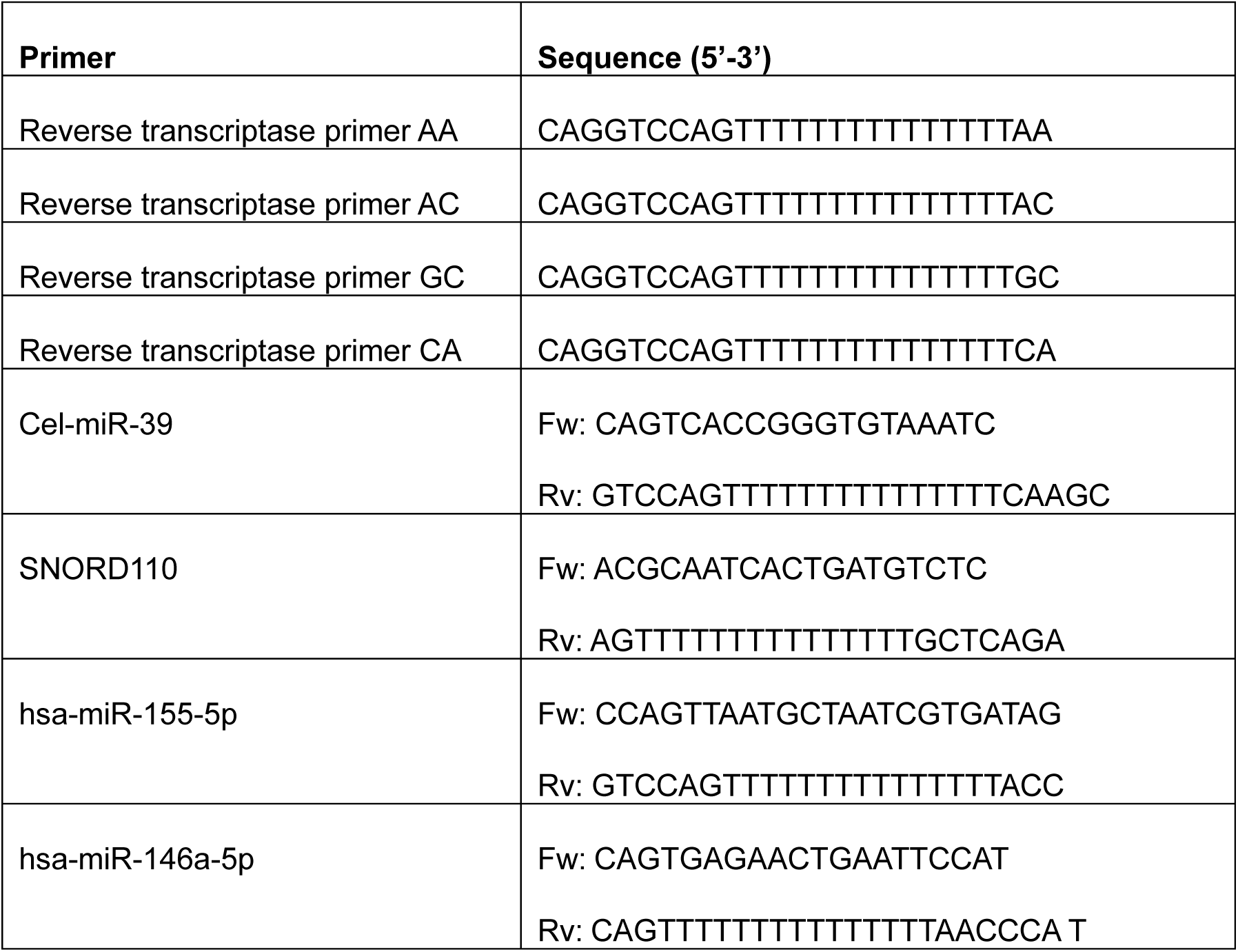
Primers and their respective sequences used for quantitative PCR analysis of miRNAs.

### mRNA and miRNA Sequencing

After total RNA extraction and quality check (see above), extracted RNAs were sent for library preparation and transcriptome analyses to GENEWIZ®. Sequencing studies were performed using a NovaSeq (Illumina) with 2x150 bp configuration and corresponding Illumina kits according to manufacturer’s instructions.

### Raw Sequence Processing and Transcript Abundance Quantification

The RNA-sequencing products were received in unprocessed fastq format. We created a genomic index using the current version of the human genome assembly (GRCH38). This index was then used for transcript abundance quantification using kallisto (Bray *et al*, 2016). We used kallisto to perform mapping/alignment of the raw reads from the reads to reference transcripts. This results in the estimated number of counts per transcript, outputted as a tabular file. The estimated count file was further processed with tximeta and tximport (Love *et al*, 2020). The implementation of tximeta facilitated loading and analyzing the data in DESeq2 (Love *et al*, 2014).

### Differential Gene Expression Analysis of mRNA of Count Data

For differential gene expression analysis, we used the tximeta package (Love *et al*, 2020) to assemble estimated count and offset matrices. This approach accounts for gene length variation across samples and retains reads mapping to homologous genes. Additionally, tximeta facilitates seamless integration with DESeq2 by providing a SummarizedExperiment object with automatic transcript annotation. The SummarizedExperiment object was converted into a DESeqDataSet for differential expression analysis. To reduce sparsity and improve computational efficiency, we filtered out low-count or zero-expression genes. Differential expression analysis was then performed using DESeq2, and significantly up- and down-regulated genes were annotated using the AnnotationDbi and org.Hs.eg.db R packages.

### Principal component analysis, hierarchical clustering, and statistical over-representation analysis

RT-qPCR data were analyzed by importing the dCt data in MATLAB, followed by normalization to the corresponding M0 condition in each case and a MinMax normalization for each of the genes across the whole dataset before running PCA (*PCAovertTimeM0normalized.m*). The explained variance was extracted and the first principal component was plotted against the corresponding time points. All curves were plotted using the plot command against time and the explained variance of the first principal component was added to the label in the y-axis.

For sequencing data, the TPM counts for all transcripts were imported in MATLAB. The script (*PCA.m*) was used for both 24 h and 72 h and a filter to remove transcripts with 3 or more empty columns (experimental conditions) was applied. The PCA command was applied, the explained variance extracted, and the first two principal components (score) were plotted, with the explained variance for each component added to the axis labels. A first individual PCA was done for each time point and a merged data set was also created to run a full PCA for all conditions across all time points.

To identify the transcript composition we downloaded the corresponding Ensembl data from biomart including Gene stable ID, Transcript Stable ID, Protein stable ID, Gene name, Gene type and Gene Synonym (Dyer *et al*, 2025). We constructed a barplot with the percentages of transcripts belonging to each category and programmed a script (*PCATranscriptsCodingvsNonCoding.m*) to run a PCA after filtering only for the protein-coding transcripts and the lncRNA transcripts. To perform our PCA we first programmed a script to collapse all the transcript contributions into a unique data point for each gene (*CreateDatasetUniqueGenes.m*). We repeated the PCA analysis for this gene-level data set either individually or as a merged data set for both time points (*PCAgeneLevel.m*). In addition, using this gene-level expression dataset we included a hierarchical analysis using the function clustergram in MATLAB, standardizing by row (experimental replicate).

For statistical over-representation analysis we uploaded the lists of selected genes in Panther database (Thomas *et al*, 2022) including only GO biological processes with p-values lower than 0.05 with the Bonferroni correction for multiple testing. The results were visualized in MATLAB for the blue and red clusters (*PlotOverRepresentationAnalysis.m*).

### Proteomics and data analysis

For protein in-gel digestion 5 µg per sample were resolved briefly in a 4-12% NuPAGE Bis-Tris gradient gel, stained with Coomassie blue and cut into small gel cubes, followed by destaining in 50% ethanol/25 mM ammonium bicarbonate. Afterwards, proteins were reduced in 10 mM DTT at 56°C and alkylated by 50 mM iodoacetamide in the dark at room temperature. Enzymatic digestion of proteins was performed using trypsin (1 µg per sample) in 50 mM TEAB (triethylammonium bicarbonate) overnight at 37°C. Following peptide extraction sequentially using 30% and 100% acetonitrile, the sample volume was reduced in a centrifugal evaporator to remove residual acetonitrile. The sample volume was filled up to 100 µl by addition of 100 mM TEAB.

Dimethyl-labelling was performed as previously reported (Boersema *et al*, 2009). Briefly, the digested samples were labelled as “Light”, “Medium” or “Heavy” by adding a combination of CH2O/NaBH3CN, CD2O/NaBH3CN or 13CD2O/NaBD3CN, respectively. Thereafter, the samples were incubated at room temperature with orbital shaking for 1 h. The labelling reaction was quenched by adding ammonia solution. After that, peptides were acidified with formic acid to reach pH ∼3. The triple-labelled samples, including a label swap, were then combined. The resultant peptide solution was purified by solid phase extraction in C18 StageTips (Rappsilber *et al*, 2003).

Before liquid chromatography tandem mass spectrometry peptides were separated on an EASY-nLC 1000 system (Thermo Scientific) via an in-house packed 30-cm analytical column (inner diameter: 75 µm; ReproSil-Pur 120 C18-AQ 1.9-µm beads, Dr. Maisch GmbH) by online reverse phase chromatography through a 225-min non-linear gradient of 1.6-32% acetonitrile with 0.1% formic acid at a nanoflow rate of 225 nl/min. Afterwards, the eluted peptides were sprayed directly by electrospray ionization into a Q Exactive Plus Orbitrap mass spectrometer (Thermo Scientific). Mass spectrometry was conducted in data-dependent acquisition mode using a top10 method with one full scan (scan range: 300 to 1,650 m/z; resolution: 70,000, target value: 3 × 10^6^, maximum injection time: 20 ms) followed by 10 fragment scans via higher energy collision dissociation (HCD; normalized collision energy: 25%, resolution: 17,500, target value: 1 × 10^5^, maximum injection time: 120 ms, isolation window: 1.8 m/z). Precursor ions of unassigned or +1 charge state were rejected. Additionally, precursor ions already isolated for fragmentation were dynamically excluded for 35 s.

Raw data files were processed in MaxQuant software (version 1.6.10.43) (Cox & Mann, 2008) using its built-in Andromeda search engine (Cox *et al*, 2011) and default settings. Spectral data were searched against a target-decoy database consisting of the forward and reverse sequences of the Homo sapiens proteome (Swiss-Prot and TrEMBL: 42,338 + 54,436 entries) downloaded on 17th January 2020 and a list of common contaminants. Corresponding labels were selected for “Light” (DimethLys0 and DimethNter0), “Medium” (DimethLys4 and DimethNter4) and “Heavy” (DimethLys8 and DimethNter8) labels. A maximum of three labelled amino acids per peptide were considered. Trypsin/P specificity was assigned.

Carbamidomethylation of cysteine was set as fixed modification. Oxidation of methionine and acetylation of the protein N-terminus were chosen as variable modifications. A maximum of two missed cleavages were tolerated. The “second peptides” search was activated. The minimum peptide length was set to 7 amino acids. False discovery rate (FDR) was set to 1% for both peptide and protein identifications.

For protein quantification, minimum ratio count of two was required. Both the unique and razor peptides were used for quantification. The “re-quantify” function was switched on. The “advanced ratio estimation” option was also chosen. Downstream data analysis was performed in R statistical environment. Reverse hits, potential contaminants and protein groups “only identified by site” were filtered out. Protein groups with at least two peptides including at least one unique peptide were retained.

Visualization of data distribution and application of the boolean criteria to select the up- and down-regulated genes for each comparison, was done using MATLAB (*ProteomicAnalysis.m*).

### Differential Gene Expression Analysis of miRNA of Count Data

The sequencing products were mapped using kallisto pseudoaligner. The resulting abundance counts were loaded in the R programming language and additional gene metadata were imported using the R/Bioconductor package tximport. The resulting gene set data structure, including the conditions to be compared were analyzed for differentially expressed genes using a pipeline based on DESeq2. A statistical significance threshold for the adjusted p-value of 0.1 was chosen, allowing relative tolerance for Type 1 error. Differentially expressed genes with an adjusted p-value equal or less than 0.1 were functionally annotated using the Bioconductor’s AnnotationDbi and selecting the human genome from the org.Hs.eg.db in order to obtain relevant information. The list of differentially expressed miRNAs was used as input for the construction of a hypothetical network of miRNA regulatory interactions with target genes including those identified by the proteomics approach.

### Network construction, simplifications and visualization in Cytoscape

To construct and simplify the initial networks we used MATLAB (*FindInteractions.m*). The inputs of the script are the list of differentially expressed genes (Degenes.mat; potential genes transitioning from one steady state to another) and our database of experimentally validated interactions (Interactions2023s.mat). The same procedure was applied to construct and simplify the 24 h and 72 h network. First, the whole database is searched for transcription factors and miRNAs reported to have a regulatory interaction on any of those differentially expressed genes. For all those edges, both sources and sinks are identified for the M1-upregulated genes and a M1 subnetwork is constructed where the total number of TFs, miRNAs, and targeted genes are identified. A similar process is performed to construct the M2 subnetwork. Afterwards, the edges of both subnetworks are included into a single list and duplicates are deleted, before performing a new identification of sources, sinks, miRNAs, TFs, and targeted genes. Then, the differentially expressed TFs are identified and the list of network edges is filtered to include only the interactions involving a differentially expressed TF, including the edges between those TFs, those TFs on miRNAs, and miRNAs on those TFs. After this first simplification, the network data is formatted for BioNetUCR and for Cytoscape for network visualization. The simplification by the differentially expressed miRNAs was performed in BioNetUCR (with the command DELETE NODES NOT IN FILE) to remove sinks and sources not participating in regulatory circuits. In parallel, we constructed a MATLAB network for the genes identified at the proteomics experiments in a similar way than for those differentially expressed genes at the transcriptomics. From this network we extracted the list of TFs and used it to filter our initial lists of differentially expressed TFs, ensuring that the final list of TFs will be composed of those which are differentially expressed, interact with miRNAs, and have at least one gene target identified at the proteomics experiment. This final simplification was performed in BioNetUCR (with the command DELETE NODES NOT IN FILE) before loading the network into Cytoscape for visualization of the topology and the means of the experimental data log2 FC(M1/M2) of the comparisons to confirm M1 upregulation (red) and M2 upregulation (blue).

### Deep search algorithm for the identification of regulatory loops

For the exploration of graphs corresponding to our networks and the identification of regulatory circuits we used MATLAB (*LoopFinder.m*). The input of the script is the list of edges of the Network. First, the stochiometric matrix was constructed for each network using an iterative routine to detect the interactions and their sign (-1,1) between all the nodes of the network. This matrix was input to the digraph function that was used to construct the graph omitting self-loops. The deep search algorithm dfsearch was implemented where the event definition included ‘edgetonew, edgetodiscovered, edgetofinished, startnode’ and the ‘edgetodiscovered’ events were highlighted. This was done recursively starting from each node of the network to find the edges ending in the same node. Within those discovered sinks we determined the shortest path (shortestpath) from each node to itself and filtered only for those interactions creating a circuit. For all the circuits the networks were filtered to keep only those including both M1 and M2 nodes. In addition, the signs of their regulation (positive or negative) were determined as the product of the signs of the individual regulations of each of the edges taking part in the respective circuit. Finally, we counted the involvement of each node across all circuits to obtain the number of positive and negative feedback loops in which they participate.

### Model construction and modifications in COPASI

Before parameter estimation, models were constructed using BioNetUCR. However, several modifications were included to ensure a proper description of bistability. BioNetUCR creates all the required reactions for all species (i.e. nodes, either mRNA or miRNA) and exports a COPASI file that includes the experimental data ready for estimating the parameters of the Ordinary Differential Equation (ODE) model. An additional set of columns for the initial concentrations of each species (genes in this case) were added to the data file output enabling the different simulated experiments to start from different initial concentrations. BioNetUCR then adds the experimental data file and the corresponding information in the COPASI file for parameter estimation. All the dynamic parameters described above are included as the fitting parameters and the data file is parsed into individual steady-state experiments in which the columns corresponding to copy numbers are included as the corresponding independent parameters in the model and the columns corresponding to expression data are included both as the corresponding independent initial concentrations of the species and the dependent variables of transient concentrations of the corresponding species at steady state.

Next BioNetUCR invokes COPASI to open the resulting file to carry out parameter estimation such that the model reproduces the provided experimental data. In COPASI, we changed the mathematical expressions of the synthesis to include Hill activation and inhibition terms to be able to capture a sigmoid-like behavior of the synthesis functions for each species. Finally, we introduced a conceptual change on the types of interactions of the model. In our previous models we used the same activation (ka) or repression (kr) parameters for all the targets of a particular species to reduce the number of parameters to be fitted (degrees of freedom). Now, to capture the individual complexity for each promoter, we included Hill terms where the half-saturation and hill coefficients could vary for each of the targets of one node. To ensure that our model simulations led to steady-states, we unchecked the “Use Newton Method” in the steady-state task.

Parameter estimation is an optimization problem to minimize the difference between simulated and experimental data (objective function), with several algorithms available in COPASI. The experimental and simulated mRNA expression data are expressed in TPM as a function of gene copy number (two for all genes) and the initial mRNA concentrations of all the species (independent variables). We developed our own fitting strategy as a pipeline of systematic parameter estimation able to search for bistability composed of an initial pipeline for individual parameter estimation and a pipeline for the progressive parameter estimation of the full model.

### Pipeline for individual parameter estimation

This pipeline is applied for each individual species to ascertain if a particular species is compatible with an overall bistable behavior of the model. Therefore, the full context (all other species) is fixed, and the fitting strategy is applied to only one species at the time.

1. Construct your model using hill expressions for synthesis functions and linear mass action kinetics for the degradation functions. Set the experiments as steady state.
2. Change the expressions of the degradation constants kd for assignments dividing the synthesis flux of the corresponding species by the initial concentration of the species (same expected at the steady state). Ensure that the compartment size is set to 1, so that synthesis and degradation can be compared. This strategy will force the optimization algorithms to identify parameters of the synthesis terms that describe a sigmoid curve (synthesis vs species concentration) intersecting a potential linear degradation curve (mass action kinetics) in at least two points, as COPASI tries to find a steady state where the synthesis is equal to the degradation.
3. Run the parameter estimation with evolutionary programming or particle swarm until finding a low objective function (in value scaling). Confirm that a bistable (or multistable) behavior is observed by changing the initial conditions (After parameter estimation on current solution statistics, update model to the corresponding experiment). For the estimation do not use Newton method (unchecked in the steady state dialog).
4. Create a new model for each species at this point and revert the expressions of the kd for fixed value returning to a linear mass action kinetics function and repeat the fitting process. Ensure that the range of Kd estimation includes the previously assigned value of Kd. Under this condition, COPASI will fine tune the synthesis parameters and determine a Kd value for a degradation function describing a linear curve crossing close to the two points previously defined by the sigmoid synthesis function for the two potential steady state concentrations defined by the experimental data.
5. Extract the steady state values and compare them to the experimental values. Confirm that a ‘bistable-like’ behavior is obtained for the current species as a function of its context. If a single steady state is found this species is not compatible with bistability for the current model and should be discarded as its behavior cannot be explained by the current model.

### Pipeline for progressive parameter estimation of the full model

The following pipeline is applied for all the species after confirmation of their compatibility with a bistable-like behavior using the individual parameter estimation pipeline. A single model is used and progressively fitted until a fully bistable model is obtained:

1. Construct your model using hill expressions for synthesis functions and linear mass action kinetics for the degradation functions. Set the experiments as steady state.
2. Change the expressions of the degradation constants kd for assignments dividing the synthesis flux of the corresponding species by the initial concentration of the species (same expected at the steady state). Ensure that the compartment size is set to 1, so that synthesis and degradation can be compared. This strategy will force the optimization algorithms to identify parameters of the synthesis terms that describe a sigmoid curve (synthesis vs species concentration) intersecting a potential linear degradation curve (mass action kinetics) in at least two points, as COPASI tries to find a steady state where the synthesis is equal to the degradation.
3. Run the parameter estimation with evolutionary programming or particle swarm until finding a low objective function (in value scaling). Confirm that a bistable (or multistable) behavior is observed by changing the initial conditions (After parameter estimation on current solution statistics, update model to the corresponding experiment). For the estimation do not use Newton method (unchecked in the steady state dialog).
4. Start reverting the expressions of the kd for fixed values one by one and repeat the fitting process. Follow the order starting with the highest degradation constants and repeat the estimation. Ensure that the range of Kd estimation includes the previously assigned value of Kd. Once a model species is well fitted, remove the corresponding parameters from the list of parameters of the parameter estimation task, so that they are not altered in posterior parameter estimations for other species.
5. Perform a fine tuning of the model parameters reentering the particular species parameters in the list of the parameter estimation task, while fixing the other species of the model (Species> type > fixed). Start with those species having the highest contribution to the objective function (Fitted Values of the Result tab of the Parameter estimation task).
6. Finally, start reverting the species type within the model to ‘reactions’ and (if necessary) repeat step 5 until reaching a full bistable model of the network.

### Steady-state simulations and parameter scans

To analyze model transitions we perturbed the initial concentrations of the individual species. To do this, a parameter scan was defined in COPASI for the object corresponding to the initial concentration of the species, for instance [CEBPA]_0 with 200 intervals, starting from a minimum of 0 to a max of two-fold the maximum concentration observed experimentally for the corresponding species. The task was set to steady state and the resulting value was plotted against the initial concentration of the corresponding species. The data was saved from the graphs and imported into MATLAB to calculate the principal component of the data over time to be plotted as a surrogate of the model status regarding the identified steady states (*PlotParameterScans.m*).

### Time-course simulations and events

All time course simulations were set to a duration beyond the transition times for all species (steady state task) and performed in COPASI using a deterministic (LSODA) method for integration. Depending on the experiments we also added specific events to simulate transient external concentrations where the trigger expression was time, the assignment target was the current concentration of the species to be evaluated, and the assignment expression was the new value given to the species. The transient concentrations of the species were plotted in COPASI against arbitrary units since the units have no meaning as the model was not fitted against time-resolved data.

### Three-dimensional parameter scan of the minimal model

For the three-dimensional parameter scan of the minimal model, first the steady-state task was run starting from an M1 or M2 experiment and the model updated to have an M1- and a M2-model. Afterwards, the Parameter Scan task was set to Time Course and the three simultaneous scans added for the initial concentrations of the species starting from 0 to the maximum experimental level observed for each of them, and 10 intervals were used for each species leading to a total of 1000 single or combined transient perturbations to the initial concentrations of the three species of interest. A report template was created to include the initial concentrations of all species, the time and the time-course data for all the species of the model. The data was imported in MATLAB and a script programmed (Plot3DscanTrajectories.m) to arrange the data, plot all the trajectories and classify them depending on the final steady state reached. Also, the script separates those trajectories according to their types according start (starting in M1 or M2) and end (ending in M1 or M2) and plot them with ‘plot3’ including either all of them or the differential trajectories.

## Acknowledgements

We thank the IMB Proteomics Core Facility for their support, especially Jasmin Cartano for her assistance with sample preparation and Mario Dejung for data analysis. Funding of the German Research Foundation supported the Q Exactive Plus system (DFG Project #240874965). The Alexander von Humboldt foundation supported R.A.M-R with the Georg Foster Fellowship (CRI 1201747 GF-E).

## Author contributions

Conceptualization, R.A.M.-R. and C.G.; Data curation, R.A.M.-R. and C.G.; Formal analysis, C.G., R.A.M-R., J.G.-C., M.A., J.T., and G.O.-B.; Funding Acquisition, A.R.-V. and R.A.M.-R.; Investigation, R.A.M.-R. and C.G.; Methodology, C.G., R.A.M-R. J.G.-C, M.A., J.T., and G.O.B.; Project administration, C.G. and A.R.-V.; Resources, C.G. and A.R.-V.; Software, R.A.M.-R. and J.G.-C.; Supervision, A.R.-V., C.G., and R.A.M.-R.; Validation, R.A.M.-R. and C.G.; Visualization, R.A.M-R and C.G.; Writing – original draft: R.A.M.-R., C.G., and A.R.-V.; Writing – review & editing, C.G., R.A.M.-R., and A.R.-V.

## Data Availability

Datasets and MATLAB-scripts are uploaded to ZENODO (DOI:10.5281/zenodo.15638350) and can be accessed via https://zenodo.org/records/15638350. Uploaded files contain:

- miRNA-Seq data
- mRNA-Seq data
- Mass spectrometry proteomics
- MATLAB-scripts

## Supplementary Figures

**Figure S1. Differential transcript expression of human macrophages under pro- and anti-inflammatory stimuli shows two different attractors for M1 and M2/M0. A.** Individual principal component analyses for the two tested time points show a higher cohesion of the clusters at 72 h. **B.** Transcript type composition indicates a majority of protein coding transcripts. **C.** Principal component analysis of the expression for protein-coding transcripts shows the two attractors clearly separated 72 h. **D.** Principal component analysis of the expression for lncRNA-coding transcripts is able to differentiate the three conditions at 24 h.

**Figure S2. Differential gene expression of human macrophages under pro- and anti-inflammatory stimuli confirms two different attractors for M1 and M2/M0. A.** Individual principal component analyses for 24 h (left) and 72 h (middle) reveal a higher cohesion of the clusters at 72 h and a combined analysis (right) indicates that M1 macrophages are still on the way to their attractor at 24 h. **B.** First top 15 GO terms according to Fold Enrichment of a statistical functional over-representation test of the M1-upregulated and the M2/M0-upregulated genes. Only GO biological processes with p-values lower than 0.05 with the Bonferroni correction for multiple testing are shown.

**Figure S3. Identification of positive and negative feedback circuits in the identified networks using a deep search algorithm starting from each node. A.** Graph of 24 h regulatory network with highlighted edges as events ending a cycle. **B.** Graph of 72 h regulatory network with highlighted edges as events ending a cycle. **C.** The level of involvement for each node is shown as the number of positive and negative feedback circuits in which each node participates.

**Figure S4. Result of individual fitting strategy to evaluate species compatibility with a bistable behavior depending on the remaining species.** Direct fitted (simulated) and experimental data comparison for the individual fitting of the species shows a good qualitative correlation and compatibility with more than one steady state as a function of the remaining model species, except for ETS2 and MIR449C. **Figure S5. Transition dependency on the modeling of other species.** The species-driven transitions were repeated upon fixation of the values of each individual remaining species to evaluate if the transition could still take place. The PCA1 of the species concentrations was used as surrogate of the system status and dashed black lines are the control transitions with all species included in the model. The green dashed lines depict the trajectories that enabled transition despite species fixation, the red ones represent those where the transition was obliterated and blue those where the transition still took place, but it is perturbed compared to control.

**Figure S6. Minimal model transitions, dependency, and hysteresic trajectories. A.** Time course simulations of subcritical (at 300-time units) and supercritical (at 4000-time units). After a subcritical transient perturbation in species concentration the system returns to the same steady state, whereas a supercritical perturbation triggers an abrupt model transition to a different steady state. The concentration of the individual species is shown in log-scale as a function of simulation time. **B.** The transition dependency on other model species shows that all species are required for at least one transition confirming the minimal model. Control transitions are shown in blue or red depending on the starting steady state. The green lines depict transitions that take place despite the fixation of species concentration and black lines depict trajectories, which transition was blocked by the fixation of the corresponding species. **C.** Differential hysteric trajectories for the same perturbations depending on the initial steady state of the system. Trajectories leading the system towards a M1 steady state are colored red and trajectories leading the system towards a M2 steady state are depicted in blue.

**Figure S7. Principal Component Analysis of human macrophages after addition of miR-155 mimic or control.** The first principal component for each independent replicate is shown over time. Cells were transfected either with miR-155 mimic (red) or control mimic (blue) at 0 h. After 4 h the pro-inflammatory stimuli were added and replaced at 24 h with the anti-inflammatory stimuli. **Figure S8. Expression of miR-146a and miR-155 in human macrophages 6, 24, 48, and 72 h after addition of M1 or M2 stimuli.** Data are shown as mean +/- SD of ddCT values normalized to unstimulated macrophages; n = 1. Higher values represent increased expression.

